# CD11c^+^ microglia promote dysfunctional T cell activation in diffuse midline glioma

**DOI:** 10.64898/2026.09.18.752751

**Authors:** Austėja Balevičiūtė, Cristina Cuesta-Marti, Jessica Hincks, Mya McCarthy Hogan, Brendan L. Sharvin, Meghan McDermott, Jessica O’Reilly, Gabriel S.S. Tofani, Aimée du Chatinier, Gerard M. Moloney, Pat Fitzgerald, Cathriona Foley, Olivia F. O’Leary, Timothy N Phoenix, Esther Hullerman, Jon D. Larson, Suzanne J. Baker, Gerard Clarke, John F. Cryan, Lily Keane

## Abstract

Paediatric diffuse midline glioma (DMG) remains refractory to immunotherapy despite T-cell infiltration, indicating that local mechanisms actively suppress anti-tumour immunity. Using complementary genetic and orthotopic mouse models, we performed lineage-resolved profiling of the DMG immune microenvironment, resolving myeloid ontogeny and microglial states by distinguishing resident microglia from infiltrating bone marrow-derived macrophages. We identified microglia as the predominant tumour-associated myeloid population and discovered selective expansion of CD11c⁺ microglia exhibiting enriched antigen-presentation and immune-regulatory transcriptional programs. Tumour-infiltrating CD4⁺ and CD8⁺ T cells exhibited chronic activation and exhaustion, while ligand–receptor analysis predicted inhibitory microglia–T cell communication. Pharmacological CSF1R inhibition depleted CD11c⁺ microglia, attenuated antigen-presentation and immune-regulatory programs, and shifted tumour-infiltrating T cells toward a less exhausted phenotype. Together, these findings identify a previously unrecognised CD11c⁺ microglia–T-cell axis that establishes immune dysfunction in DMG and provide a rationale for combining myeloid- and T cell-targeted immunotherapies.

## Main

Paediatric diffuse midline glioma (DMG) characterised by the H3K27M mutation and formerly known as diffuse intrinsic pontine glioma (DIPG), remains one of the most aggressive childhood brain tumours. DMG arises along midline structures such as the pons, thalamus and spinal cord^1^. The disease typically presents in early childhood, with a median age at diagnosis of seven years and a median survival of less than one year^2^. Due to its diffuse and infiltrative nature within critical brainstem structures, surgical resection is not feasible^3^, and treatment is largely limited to fractionated radiotherapy, which provides only transient symptomatic relief and modest survival benefit^4^.

The discovery of the recurrent H3K27M mutation, substituting lysine 27 of histone H3 with methionine, significantly advanced our understanding of DMG biology^1,5^. This mutation disrupts the activity of the polycomb repressor complex 2 (PRC2), resulting in global reduction of H3K27 trimethylation and widespread epigenetic dysregulation^6^. However, H3K27M alone is insufficient to drive tumorigenesis, which require additional oncogenic alterations such as platelet derived growth factor beta (PDGFB) signalling^7^. These important molecular insights have not as yet translated to effective therapies.

Recent clinical efforts have sought to enhance antitumour immunity in DMG through strategies that either increase T cell activity or relieve inhibitory immune regulation. Adoptive cell therapies, including CAR-T cells targeting tumour-associated antigens such as disialoganglioside (GD2)^8^ or B7-family immunoregulatory protein B7-H3^9^, demonstrate that T cells can access and recognise DMG tumours but durable antitumour responses remain rare, suggesting active constraints imposed by the tumour microenvironment (TME)^10^. Similarly data from adult glioblastoma and paediatric high-grade glioma studies, demonstrate T cells frequently exhibit dysfunctional or exhausted phenotypes, characterised by impaired effector function and sustained expression of inhibitory receptors^10–15^. Therefore, strategies to enhance endogenous T-cell responses in glioblastoma have largely focused on increasing T-cell activity either by: (i) relieving inhibitory signalling through immune checkpoint blockade, including Programmed Cell Death Protein 1 (PD-1) and Cytotoxic T-Lymphocyte–Associated Protein 4 (CTLA-4)^16–19^, or (ii) by providing activating signals through co-stimulatory receptors such as Tumor Necrosis Factor Receptor Superfamily Member 9 (4-1BB/TNFRSF9) and Glucocorticoid-Induced TNFR-Related Protein (GITR/TNFRSF18)^20,21^. However, these approaches have failed to overcome the profound immunosuppressive nature of the glioma TME, remain constrained by safety limitations associated with systemic immune activation and provide only modest clinical benefits to patients^20–24^.

Despite limited direct evidence in DMG, studies in adult and paediatric high-grade gliomas suggest that T-cell function is constrained within an immunosuppressive TME, in which tumour-associated myeloid cells (TAMs) act as key regulators alongside other intrinsic and extrinsic factors^11,12,25,26^. In adult glioblastoma, TAMs comprise both resident microglia and infiltrating bone marrow-derived macrophages (BMDMs), which differ in ontogeny and function^27–29^. However, in DMG, these populations have often been analysed as a single compartment, limiting insight into their distinct contributions to immune regulation^27,29–31^. This lack of resolution may obscure lineage-specific mechanisms of T cell dysfunction within the TME. Recent advances have established that microglia and BMDMs can be reliably distinguished using lineage markers such as purinergic receptor 12 (P2RY12) and integrin alpha 4 (ITGA4/CD49d) respectively^32^, enabling more precise dissection of their roles in shaping T cell function within the DMG TME that remains to be investigated.

Among broadly described TAMs, emerging evidence indicates that microglia play an active role in DMG progression^25,28,33,34^. Brainstem microglia have been shown to enhance tumour growth compared to cortical microglia^35^. Both experimental and clinical studies demonstrate that microglia within the DMG microenvironment adopt pro-tumorigenic phenotypes that promote tumour growth and invasion^30,36,37^. Similarly, we previously showed that microglia can be epigenetically reprogrammed toward anti-tumour states, highlighting their therapeutic potential^36,37^. Microglia exhibit substantial functional heterogeneity, adopting context-dependent activation states in response to local cues^38–40^. Among these, CD11c^+^ microglia, defined by expression of ITGAX, represent a dynamic activation state associated with phagocytosis, immune modulation, and antigen presentation across development and disease^41–45^. These cells have been implicated in developmental myelination and inflammatory responses in the cerebellum^45^. Recently we showed that CD11c^+^ microglia are the dominant population in human DMG and that their transcriptomic signatures overlap with microglia in the developing brainstem^46^. However, their functional role within the DMG TME remains unknown.

Here, we sought to define the immune mechanisms underlying T cell dysfunction in DMG by first characterising T cell activation states within the TME and then determining how distinct tumour-associated myeloid populations may shape these responses. Importantly many preclinical immunotherapy studies to date have been conducted in xenograft models using human-derived DMG cells where the potential immunosuppressive nature of the TME is missing^47^. Here, we performed a lineage-resolved profiling of the DMG TME in two cutting edge immunocompetent mouse models, distinguishing resident microglia from infiltrating BMDMs using established ontogeny markers and further interrogating microglial heterogeneity through analysis of a CD11c^+^ microglial activation state. We show that tumour-infiltrating T cells exhibit features of chronic activation and exhaustion, identify microglia as the dominant myeloid population associated with inhibitory signalling, and reveal that microglia–T cell interactions are enriched for checkpoint-mediated pathways in the absence of canonical co-stimulatory signalling. Together, these findings define a distinct microglia–T cell axis in DMG and highlight myeloid ontogeny and the microglial state as critical determinants of immunotherapy resistance, informing strategies to improve treatment response.

## Results

### T cells infiltrate DMG tumours but exhibit an exhaustion-like dysfunctional activation state

Despite current advancements in CAR-T cell research and robust expression of tumour specific CAR-T cell antigens such as GD2, cell-based therapies show limited efficiency in the treatment of DMG^8,47^. We therefore hypothesized that T cells acquire activation states within the DMG TME that may prevent them from efficiently engaging and eliminating DMG tumour cells (**Fig. 1**). To this end, we employed two complementary immunocompetent murine models of DMG: (i) an orthotopic transplantation model, in which tumour cells, derived from in utero electroporation introducing *H3f3a^K27M^*, *Pdgfra^D842V^* and dominant-negative *Trp53* (DNp53), were implanted into the pons of adult C57BL/6 mice (500,000 cells; tumours are lethal within approximately 50–60 days^48,49^) here-after referred to as O-DMG (**Fig. 1A**), and (ii) a tamoxifen-inducible *Nestin-CreER^T^*^2^ genetically engineered model carrying *H3f3a^K27M^*, *Trp53* loss, and *PDGFRA^V544ins^* (tumours are lethal within approximately 90–100 days) here-after referred to as G-DMG^2^.

**Fig. 1:**
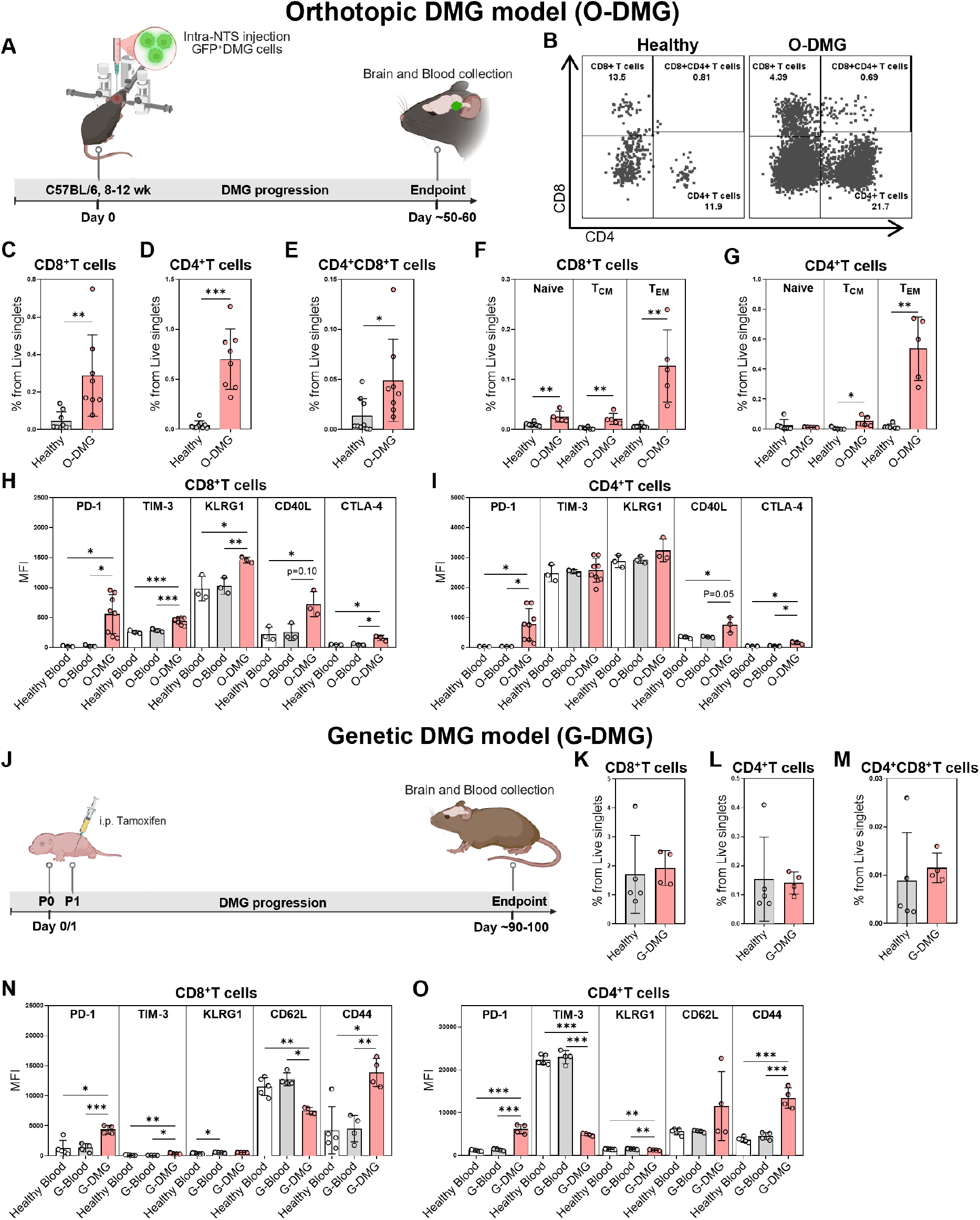
T cells infiltrate DMG tumours and exhibit an exhausted and dysfunctional activation state. **A)** Experimental design of the orthotopic immunocompetent DMG murine model (A-I); NTS – nucleus tractus solitarius. T cell phenotyping by flow cytometry: **B**) gating strategy and percentages of healthy midbrain/hindbrain (healthy) and DMG tumour (O-DMG)-associated CD4^+^, CD4^+^CD8^+^, and CD8^+^ T cells from CD45^+^ immune cells. The percentage of immune populations in healthy HB/MB and O-DMG from live singlets: **C**) CD8^+^ T cells; **D**) CD4^+^ T cells; **E**) CD4^+^CD8^+^ T cells. Naive (CD62L^+^CD44^-^), central-memory (T_CM_; CD62L^+^CD44^+^) and effector-memory (T_EM_; CD62L^-^CD44^+^) subpopulations of: **F**) CD8^+^ T cells; **G**) CD4^+^ T cells. Median fluorescence intensity (MFI) of PD-1, TIM-3, KLRG1, CD40L and CTLA-4 from: **H**) CD8^+^ T cells; **I**) CD4^+^ T cells. Healthy blood and O-DMG mice blood (O-Blood) T cells were used as controls for MFI estimation. **J**) Experimental design of the genetic immunocompetent murine DMG model (J-O). The percentage of immune populations in healthy brain versus G-DMG tumour from live singlets: **K**) CD8^+^ T cells; **L**) CD4^+^ T cells; **M**) CD4^+^CD8^+^ T cells. MFI of PD-1, TIM-3, KLRG1, CD62L and CD44 from: **N**) CD8^+^ T cells; **O**) CD4^+^ T cells. Healthy non-induced and G-DMG mice blood (G-Blood) T cells were used as controls for MFI estimation. i.p. - intraperitoneal injection; n=3-8/group; ns – non-significant; *p<0.05; **p<0.005; ***p<0.001.

To investigate whether and to what extent T cells infiltrate O-DMG tumours, we first performed flow cytometry analysis of T cell subsets (**Supplementary 1A-B**). In the orthotopic murine model, as 100% of tumours occur in the brainstem, we compared O-DMG tumours to the corresponding healthy midbrain/hindbrain (MB/HB) regions, referred to as “healthy” (**Fig. 1B, Supplementary 2A**). Flow cytometry analysis revealed that in comparison to the healthy brain tissue, O-DMG tumours exhibit a significant increase in CD8^+^ (**Fig. 1C**), CD4^+^ (**Fig. 1D**) and exhaustion-associated double-positive CD4^+^CD8^+^ T cells (**Fig. 1E**), where an observed higher CD4/CD8 ratio is consistent with a relatively bigger population of CD4^+^ compared to CD8 ^+^ T cells within the TME (**Supplementary S2B**). However, despite a significant increase observed in the percentage of O-DMG-associated T cell populations, the T cells represent a minor population as they only make up to 1% of TME (**Fig. 1C-E**). To further assess O-DMG-associated T cell functional and memory status, T cells from O-DMG and healthy mouse peripheral blood were used for comparison. Upon entry to the tumour, majority of O-DMG-associated CD8^+^ and CD4^+^ T cells were effectors (**Fig. 1F, 1G**). We therefore carried out further activation and cytotoxicity phenotyping. Indeed, we found that O-DMG-associated T cells exhibited dysfunctional activated phenotype marked by the upregulation of PD-1 on the effector CD8^+^ (**Fig. 1H**) and CD4^+^ T cell subsets (**Fig. 1I**). Additionally, O-DMG-associated CD8^+^ T cells had upregulated inhibitory receptors TIM-3 and CTLA-4, terminal differentiation/exhaustion-associated marker KLRG1 and costimulatory ligand CD40L compared to healthy or O-DMG peripheral blood counterparts (**Fig. 1H**). O-DMG-associated CD4^+^ T cells shared some of this dysfunctional phenotype with CD8^+^ T cells by upregulating CD40L and CTLA-4 (**Fig. 1I**). Moreover, O-DMG-infiltrating CD8^+^ and CD4^+^ T cells exhibited significantly increased expression of proliferation-associated Ki-67, cytotoxic granzyme B, exhaustion-associated CD39 compared to the T cells in the lymph nodes of O-DMG tumour-bearing mice (**Supplementary 2C-D**), suggesting expanded partially dysfunctional T cell activation state upon entry to the DMG tumour.

Based on observed dysfunctional O-DMG-associated T cell phenotype, we have validated these findings in the G-DMG model (**Fig. 1J**). Given that 10-15% of G-DMG tumours occur in other brain regions, we compared whole G-DMG brain to the healthy whole brain from non-induced animals as controls. Compared to O-DMG, in G-DMG tumours we did not observe an increase in T cell infiltration compared to healthy noninduced controls, possibly due to the whole brain comparison (**Fig. 1K-M**). In contrast to O-DMG, the G-DMG model showed a lower CD4/CD8 ratio, indicating CD8^+^ T cells a dominant population compared to CD4^+^ T cells (**Supplementary 2B**). However, despite these differences observed in both DMG models, further phenotyping of T cell functional and memory status revealed that similarly to O-DMG, the majority of G-DMG-associated CD8^+^ (**Fig. 1N**) and CD4^+^ T cells (**Fig. 1O**) adopted effector phenotype. Moreover, similarly to O-DMG, G-DMG-associated T cells displayed an increased expression of CD44 and decreased expression of lymph node homing receptor CD62L compared to circulating blood T cells, suggesting antigen experience and acquisition of effector-like phenotype upon entry into the brain tumour microenvironment, accompanied by loss of a naïve T cell state (**Fig. 1N-O**). Consistently with O-DMG model, G-DMG-associated T cells exhibited dysfunctional activated phenotype marked by the upregulation of PD-1 on effector CD4^+^ (**Fig. 1N**) and CD8^+^ (**Fig. 1O**) T cells subsets. Additionally, as observed in O-DMG, G-DMG-associated CD8^+^ T cells had increased expression of inhibitory receptor TIM3 (**Fig. 1O**). Together, findings from both O-DMG and G-DMG support a model in which DMG TME drives CD8⁺ T cells and CD4^+^ T cells into a chronically stimulated, partially functional state associated with emerging exhaustion-like dysfunction.

Next, we FACS-sorted CD4^+^ and CD8^+^ T cells from O-DMG tumours and corresponding healthy MB/HB controls for bulk transcriptomics to further characterize tumour-associated T cell activation states (**Fig. 2; Supplementary 2E**). Bulk RNA-sequencing analysis investigated functional and transcriptional gene signatures as well as the exhaustion gene module score, reflecting the overall expression of the exhaustion gene signature. Similarly to flow cytometry analysis, RNA sequencing revealed that CD8^+^ (**Fig. 2A**) and CD4^+^ T cells (**Fig. 2B**) downregulate genes associated with naive T cell state and acquire activation/chronic stimulation and terminal exhaustion related phenotypes but retain some cytotoxicity (**Fig. 2C-F**). CD8^+^ T cells display an exhaustion-like dysfunctional state (**Fig. 2C-D; Supplementary 3A**) with atypical expression of suppression gene signatures such as significant elevation of *Foxp3, Nrp1, Il10* and a trend towards elevated *Ccr8* expression (**Supplementary 3B**). Similarly, but to a greater extent, CD4^+^ T cells show gene signatures related to exhaustion (**Supplementary 3C**) and inhibitory signalling (**Supplementary 3D**). Interestingly, Enrichr enrichment analysis using TRRUST^50^ transcription factor database mining revealed that both, O-DMG-associated CD8^+^ (**Supplementary 4A-B**) and CD4^+^ T cells (**Supplementary 4C-D**) displayed activation, exhaustion and stress response-related transcriptional signatures, suggesting that both T cell subsets adopt a dysfunctional activation state. Gene Ontology (GO) Molecular Function terms analysis further revealed O-DMG-associated CD8^+^ T cells displayed rewired Nuclear Factor or Activated T cells (NFAT) signalling axis that suggests favoured exhaustion/tolerance over productive effector function (**Supplementary 4E-F**). Similarly, CD4^+^ T cells displayed gene programs related to regulatory-like T cell phenotypes and weakened stress response, possibly leading to the loss of transcriptional regulation and functional responsiveness (**Supplementary 4G-H**). Moreover, Metascape enrichment analysis of both CD8^+^ (**Supplementary 5A-B**) and CD4^+^ T cells (**Supplementary 6A-B**) revealed upregulation of immune and inflammatory pathways, including interferon and cytokine signalling, together with downregulation of ribosome biogenesis, transcription/translation-associated processes, and T-cell receptor signalling. Collectively, these findings suggest that the DMG tumour microenvironment promotes a regulatory-like, chronically stimulated, partially functional, and exhaustion-prone T-effector state.

**Fig. 2:**
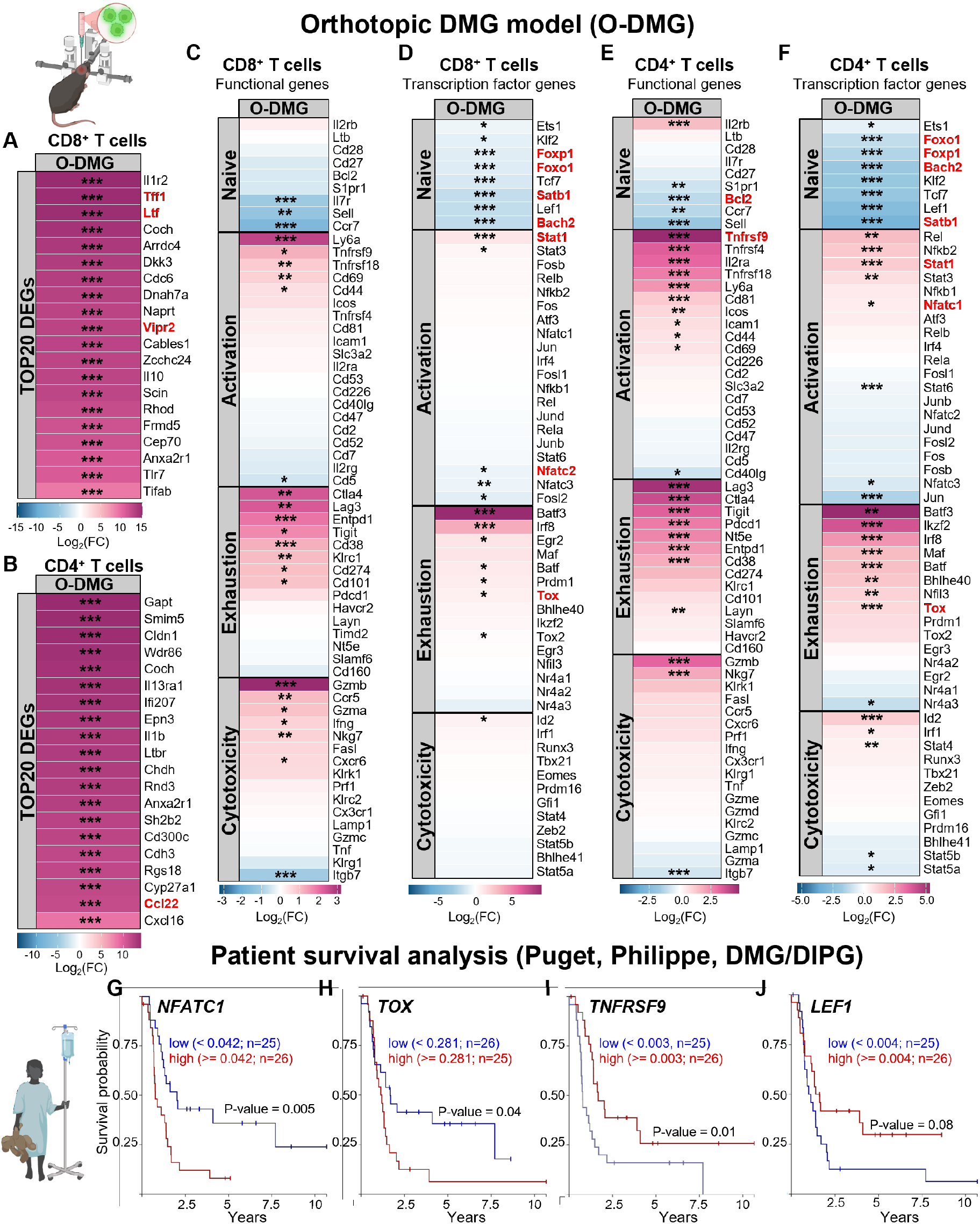
DMG-associated T cells display exhaustion and regulatory-like transcriptional programs related to dysfunction. Bulk RNA-sequencing of CD8^+^ T cells (A-G) and CD4^+^ T cells (H-N) sorted from the orthotopic diffuse midline glioma (O-DMG) tumour region and the corresponding healthy midbrain/hindbrain (MB/HB) region used as a non-tumour control. Top 20 differentially expressed genes (DEGs) in: **A)** CD8^+^ T cells and **B**) CD4^+^ T cells. CD8^+^ T cell naïve, activation, exhaustion and cytotoxicity associated **C**) functional and **D**) transcription factor genes visualized as an expression heatmap. CD4^+^ T cell naïve, activation, exhaustion and cytotoxicity associated **E**) functional and **F**) transcription factor genes visualized as an expression heatmap. Significance: *p<0.05; **p<0.005; ***p<0.001, otherwise non-significant; n=3-4/group. Red coloured genes in the A-F heatmaps were investigated in Puget et al. (2012) patient cohorts for the survival analysis; Kaplan–Meier survival curves for Puget et al. (2012) DIPG/DMG patients stratified by median expression of indicated genes (high vs. low), visualized using UCSC Xena: **G**) NFATC1; **H**) TOX; **I**) TNFRSF9; **J**) LEF1.

To determine whether the T cell transcriptional programmes identified in our orthotopic and genetic DMG models were clinically relevant, we compared our findings with the expression of these genes in DMG patient cohorts^51^. Only significantly altered genes in the O-DMG tumour-associated T cells were selected (marked in red font; **Fig. 2A-F**) and tested to see if their expression had an impact on patient survival (**Fig. 2G-J**). Elevated expression of *NFATC1* (**Fig. 2G**), a marker of sustained T-cell receptor (TCR) signalling, and the exhaustion-associated transcription factor *TOX* (**Fig. 2H**) were associated with poorer patient survival, consistent with chronic T-cell activation and progressive dysfunction. Conversely, reduced expression of *LEF1* (**Fig. 2I**), which maintains naïve and memory T-cell identity, and the co-stimulatory receptor *TNFRSF9* (**Fig. 2J**) also correlated with adverse clinical outcomes, suggesting impaired T-cell persistence and activation in patients with DMG. Consistent with these clinical findings, our data show that murine tumour-infiltrating T cells, particularly CD4⁺ T cells, displayed increased expression of *Nfatc1* and *Tox*, together with reduced *Lef1* and *Tnfrsf9* (**Fig. 2C-F**). These findings support a model in which chronic activation and inhibitory/exhaustion-associated signalling programmes in CD4⁺ and CD8⁺ T lymphocytes drive progressive T-cell dysfunction, limiting their capacity to mount an effective anti-tumour immune response within the DMG microenvironment.

### DMG-associated TAMs exhibit a distinct immunosuppressive activation state

Given the presence of chronically activated/exhausted-like dysfunctional CD4^+^ and CD8^+^ T cells in DMG tumours, we next investigated TAM populations as potential mediators of immune dysfunction (**Fig. 3**). Here we compared O-DMG tumour with corresponding healthy MB/HB, focusing on microglia (P2RY12^+^, CD49d^-^), myeloid-derived suppressor cells (MDSCs; P2ry12^int^, CD49d^int^) and bone-marrow derived macrophages (BMDMs; P2RY12^-^, CD49d^+^; **Fig. 3A, Supplementary 7A**). O-DMG-tumours showed an increase in the percentage of CD11b^+^ myeloid population (**Supplementary 7B**). CD11b^+^ myeloid cells were further distinguished into BMDMs, MDSCs and microglia. An increasing trend in BMDMs infiltration was observed in O-DMG tumours (**Fig. 3B**), but no changes were observed in the abundance of MDSCs (**Fig. 3C**) or microglia (**Fig. 3D**). Despite the infiltration of BMDMs into the tumour, they only took up to 10-15% of the CD11b^+^ compartment, similarly to MDSCs (**Fig. 3E**). Whereas, microglia remained the predominant myeloid population, comprising up to 70% of CD11b^+^ myeloid cells in the O-DMG tumour (**Fig. 3E**). Further phenotyping revealed no significant differences in the expression of inhibitory or activating molecules, where CD11c expression on O-DMG-associated global microglia was higher but did not reach statistical significance (**Fig. 3F**). Despite this, given that CD11c^+^ microglial states have previously been reported in developmental, neurodegenerative, and glioblastoma contexts^41–45,52^, coinciding with the developmental emergence of DMG, we hypothesized that this activation state might play a role in DMG progression, potentially aiding in dysfunctional T cell immune response. We therefore examined the presence of CD11c⁺ microglia (**Fig. 3G**), that we previously described as the predominant microglia subtype within the healthy developing brainstem and human DMG^46^. Here, O-DMG tumours exhibited a significant expansion of the CD11c⁺ microglial subpopulation compared to the healthy counterparts (**Fig. 3H**). This subset comprised approximately 15-20% of total microglia within the DMG microenvironment (**Fig. 3I**). Notably, CD11c⁺ microglia displayed increased expression of the inhibitory ligand PD-L1 (**Fig. 3J**) and toll-like receptor 2 (TLR2) as well as dysregulated expression of MHC-II, CD86, CD40 and CLEC7A (**Supplementary 7C**).

**Fig. 3:**
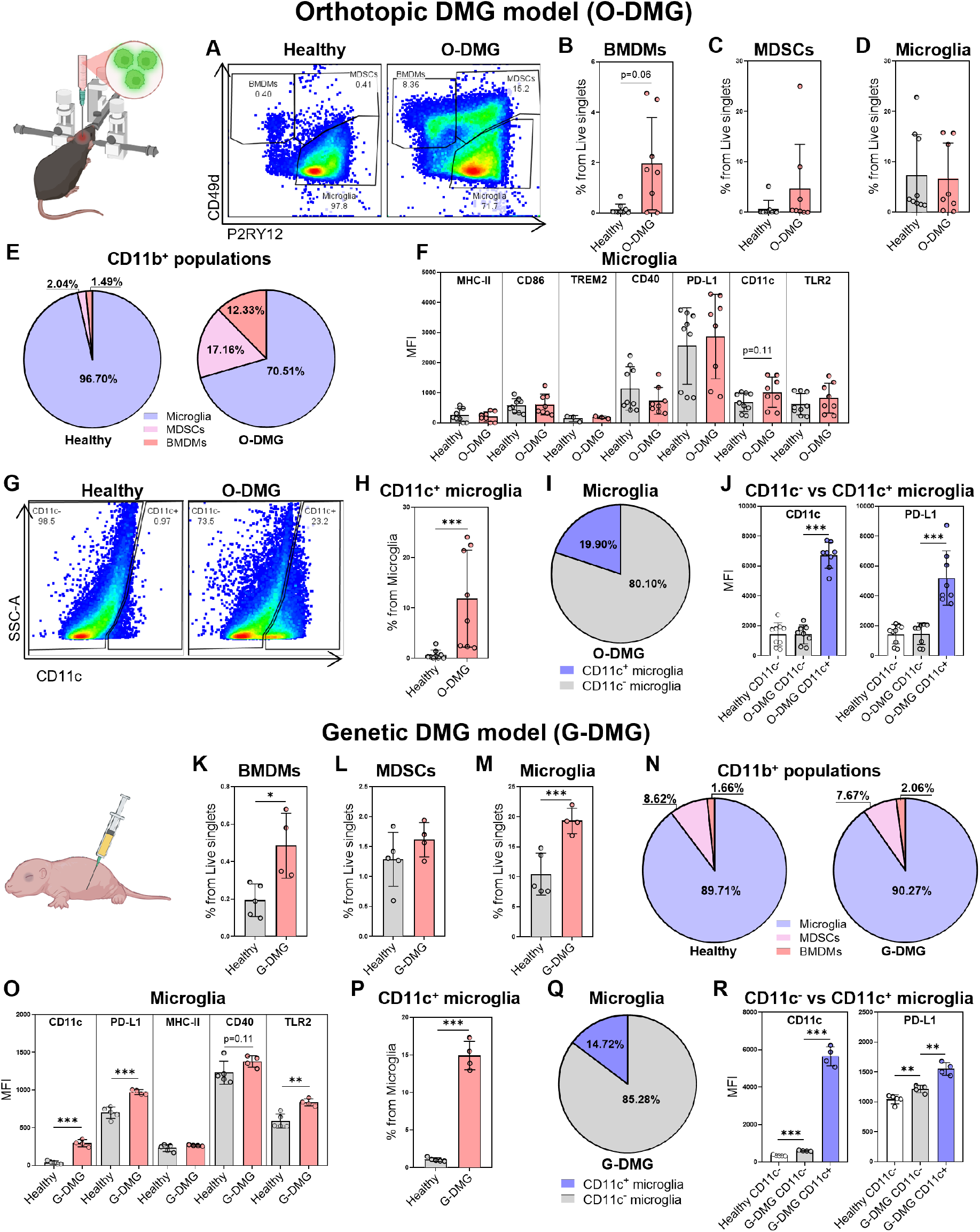
TAM populations acquire immunosuppressive activation states in DMG. TAM phenotyping of orthotopic DMG (O-DMG; A-J) and genetic DMG (G-DMG; K-R) by flow cytometry. **A**) Gating strategy of macrophages, myeloid derived suppressor cells (MDSCs) and microglia in healthy hindbrain/midbrain (HB/MB) and corresponding DMG tumour region. The percentage of immune populations in healthy HB/MB and O-DMG from live singlets: **B**) bone marrow-derived macrophages (BMDMs); **C**) Myeloid-derived suppressor cells (MDSCs); **D**) Microglia. **E**) Venn diagram showing percentages of microglia, BMDMs and MDSCs from CD11b^+^ myeloid cells in O-DMG. **F**) Median fluorescence intensity (MFI) of MHC-II, CD86, TREM2, CD40, PD-L1 and CD11c from global microglia population. **G)** Gating strategy of CD11c^+^ and CD11c^-^ microglia in healthy forebrain (FB) and corresponding DMG non-tumour region (NTR) with healthy midbrain/hindbrain (MB/HB) and its corresponding DMG tumour. **H**) Percentages of CD11c^+^ microglia from global microglia population; **I**) Venn diagram showing the percentage of CD11c^+^ microglia from global microglia population. **J**) MFI of CD11c and PD-L1 on CD11c^+^ and CD11c^-^ microglia in healthy MB/HB, NTR and O-DMG tumour. The percentage of immune populations from live singlets in healthy brain and G-DMG tumour: **K**) Macrophages; **L**) Myeloid-derived suppressor cells (MDSCs); **M**) Microglia. **N**) Venn diagram showing percentages of microglia, macrophages and MDSCs from CD11b^+^ myeloid cells in G-DMG. **O**) Median fluorescence intensity (MFI) of CD11c, PD-L1, MHC-II, CD40 and TLR2 from global microglia population. **P**) Percentages of CD11c^+^ microglia from global microglia population. **Q**) Venn diagram showing the percentage of CD11c^+^ microglia from global microglia population. **R**) MFI of CD11c and PD-L1 on CD11c^+^ and CD11c^-^ microglia in healthy MB/HB, NTR and G-DMG tumour. n=3-8/group. Significance: *p<0.05; **p<0.005; ***p<0.001, otherwise non-significant.

In the genetic G-DMG model, myeloid cell populations were assessed comparing whole non-induced brains versus G-DMG brains. Similarly to O-DMG, in the G-DMG tumours, we found an increase in the percentage of CD11b^+^ myeloid cells (**Supplementary 7D**). Moreover, similarly to O-DMG, G-DMG tumours were significantly infiltrated by peripheral BMDMs (**Fig. 3K**), but no changes were observed in MDSCs (**Fig. 3L**). Despite no changes in O-DMG tumours, the percentage of G-DMG-associated microglia was significantly increased in G-DMG brains (**Fig. 3M**), where they remained the predominant myeloid population, comprising up to 90% CD11b^+^ myeloid cells (**Fig. 3N**). Unlike O-DMG, G-DMG-associated global microglia population showed a pronounced upregulation of PD-L1 (**Fig. 3O**) and TLR2, consistent with an activated and potentially immunoregulatory state. However, similarly to O-DMG, but to a greater extent, G-DMG-associated global microglia displayed a significant increase in CD11c expression (**Fig. 3O**). Similarly to O-DMG, G-DMG tumours exhibited a significant expansion of the CD11c⁺ microglial subpopulation compared to the healthy counterparts (**Fig. 3P**). This subset comprised approximately 15% of total microglia within the DMG microenvironment (**Fig. 3Q**). Notably, G-DMG-associated CD11c⁺ microglia displayed significantly increased expression of the inhibitory ligand PD-L1 (**Fig. 3R**), as well as TLR2 and dysregulation of MHC-II, CD86, CD40 and CLEC7A (**Supplementary 7E**).

Together, these findings suggest that CD11c^+^ microglial activation state emerges in the DMG tumours and possibly contributes to the immunosuppressive microenvironment. We have shown that CD11c^+^ microglia have an increased expression of inhibitory checkpoint molecule PD-L1 and activation-associated TLR2, together with altered regulation of antigen presentation and co-stimulatory molecules, suggesting that this microglial activation state can participate in driving T cell dysfunction.

### CD11c^+^ microglia represent an immunomodulatory activation state associated with T cell dysfunction in DMG

Following the identification of a CD11c^+^ microglial activation state in DMG, we aimed to further validate and functionally characterize the microglial compartment within the DMG microenvironment. Therefore, we next performed bulk RNA sequencing of sorted O-DMG tumour-associated microglia comparing them with the healthy counterparts (**Fig. 4**). Transcriptomics data revealed that O-DMG tumour-associated microglia upregulated immunosuppression, tumour support, hypoxia and metabolically dysfunction-associated genes (**Fig. 4A-B**). The global microglial population had a significant upregulation of CD11c^+^ microglial signature-associated genes, confirming the presence of CD11c^+^ microglial activation state (**Fig. 4C**). Moreover, global microglia had a prominent antigen presentation signature, suggesting possible interactions with adaptive immunity (**Fig. 4D**). Cell-cell sender-receiver predicted interaction analysis has revealed strong potential interactions from microglia signalling towards CD8^+^ T cells and to a greater extent towards CD4^+^ T cells (**Fig. 4E**). Consistent with this, TTRUST (**Supplementary 8A-B**), GO Molecular Function (**Supplementary 8C-D**) and Metascape network analyses (**Supplementary 8E-F**) revealed O-DMG-associated microglia exhibited activated, interferon-driven inflammatory and proliferative state while suppressing homeostatic programs, indicative of an activated immune signalling state potentially involved in T cell communication. Supporting this observation, subsequent ligand–receptor interaction analyses identified multiple putative immune regulatory interactions between microglia and T cells, including inhibitory PD-L1–PD-1, LAG3-MHC-I, CD200-CD200R4, SLAMF receptor family interactions and non-canonical co-stimulatory interactions such as LTBR–LIGHT, CD30–CD30L and KLKR1-RAET1A signalling axes (**Fig. 4F-G**). However, canonical co-stimulatory CD28-CD80/CD86, ICOS-ICOSL, CD40-CD40L signalling axes were absent (**Fig. 4F-G**), supporting the hypothesis by which chronic antigen exposure with dysfunctional co-stimulation and present inhibitory signalling could lead to T cell suppression, and ultimately, exhaustion.

**Fig. 4:**
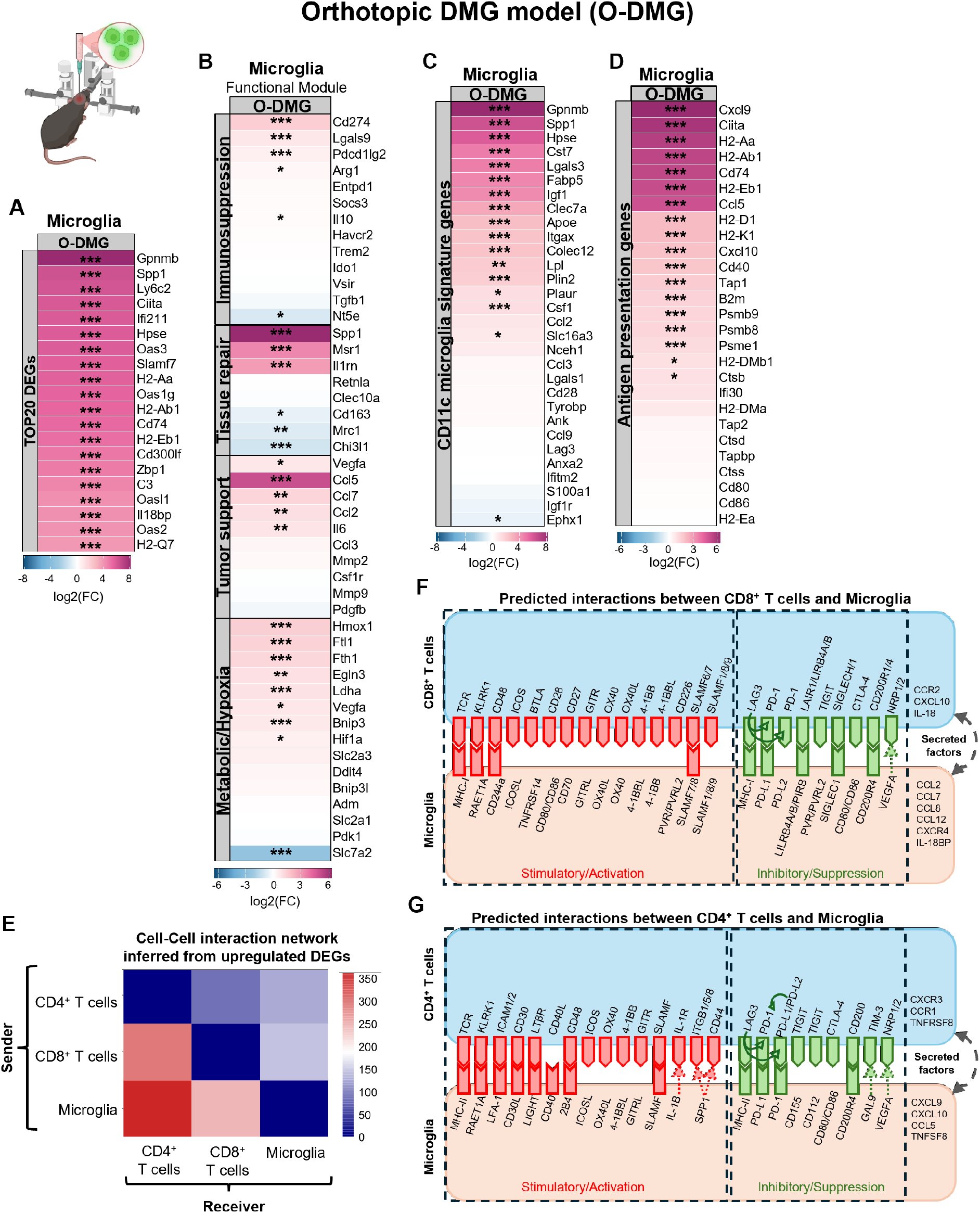
Inferred ligand–receptor interactions between T cells and microglia, including CD11c^+^ microglia activation state. Bulk RNA-sequencing of microglia (CD11b^+^CD49^-^P2RY12^+^) in the O-DMG tumour region versus healthy midbrain/hindbrain region: **A**) TOP20 differentially expressed genes (DEGs); **B**) Immunosuppression, tissue repair, tumour support and metabolic/hypoxia-associated gene heatmap. **C**) CD11c microglial gene signature heatmap. **D**) Antigen presentation gene signature heatmap. **E**) Cell-cell interaction network generated as sender-receiver interaction pairs inferred from bulk RNA sequencing data from sorted O-DMG associated versus healthy microglia, CD4^+^ and CD8^+^ T cells. Inferred ligand-receptor pairs between microglia and: **F**) CD4^+^ T cells; **G**) CD8^+^ T cells. Inferred ligand-receptor interactions based on DEGs, using CellTalkDB, CellPhoneDB and OmniPath. n=4/group. Significance: *p<0.05; **p<0.005; ***p<0.001, otherwise non-significant.

Interestingly, microglia appear to upregulate genes related to chemokine signalling and T cell recruitment, such as *Cxcl9*, *Cxcl10,* and *Ccl2* with receptor encoding gene *Ccr2* upregulated in activated CD8^+^ T cells and *Cxcr3* in CD4^+^ T cells (**Fig. 4F-G**). Therefore, to investigate whether DMG alters systemic immune signalling, we measured circulating cytokines in blood plasma of the O-DMG-bearing mice (**Supplementary 9**). We observed a trend towards increased CCL2 (**Supplementary 9N**), suggesting altered peripheral inflammatory signalling and immune-cell trafficking. Moreover, O-DMG-bearing mice exhibited significantly reduced CX3CL1 (**Supplementary 9S**) and a trend towards increased IL-27 (**Supplementary 9K**), consistent with an immunomodulatory environment. These findings suggest that tumour-associated microglia possibly contribute may primarily regulate the behaviour of the sparse T cells present within the tumour, influencing their retention, positioning, or local interactions rather than promoting effective T cell accumulation or activation. Together, these data support the existence of coordinated microglia–T cell crosstalk in DMG while highlighting candidate chemokine pathways that may contribute to the restricted immune landscape of these tumours.

Together, these findings identify a CD11c⁺ microglia state enriched in DMG tumours that exhibits features consistent with immunomodulatory signalling. The absence of canonical co-stimulatory pathways, combined with enrichment of checkpoint interactions, suggests that microglia, particularly CD11c^+^ microglial state, may engage T cells in a manner that favours dysfunctional activation rather than effective anti-tumour responses. Given that microglia are poor antigen-presenting cells, at least compared to dendritic cells, within the CNS immune-privileged environment^53^, these results suggest they may limit full T cell activation.

### CSF1R inhibition, using PLX3397, reduces CD11c^+^ microglial signatures and reshapes the myeloid landscape in DMG

Based on our predicted receptor–ligand interaction analysis (**Fig. 4**), we next investigated mechanisms underlying T-cell suppression and exhaustion within the DMG microenvironment. We hypothesized that tumour-associated myeloid cells play a central role in driving T-cell dysfunction, and that perturbing these populations could reshape T-cell states. To test this, we targeted colony-stimulating factor 1 receptor (CSF1R) signalling using pexidartinib (PLX3397), a potent small-molecule inhibitor of CSF1R (**Fig. 5**). CSF1R is expressed predominantly on circulating monocytes, macrophages, and microglia, where it regulates survival, proliferation, and differentiation^54^. In tumours, CSF1R signalling supports the accumulation and maintenance of TAMs, key mediators of immunosuppression^55,56^, which promote T-cell dysfunction through inhibitory ligands and cytokines^57^.

**Fig. 5.**
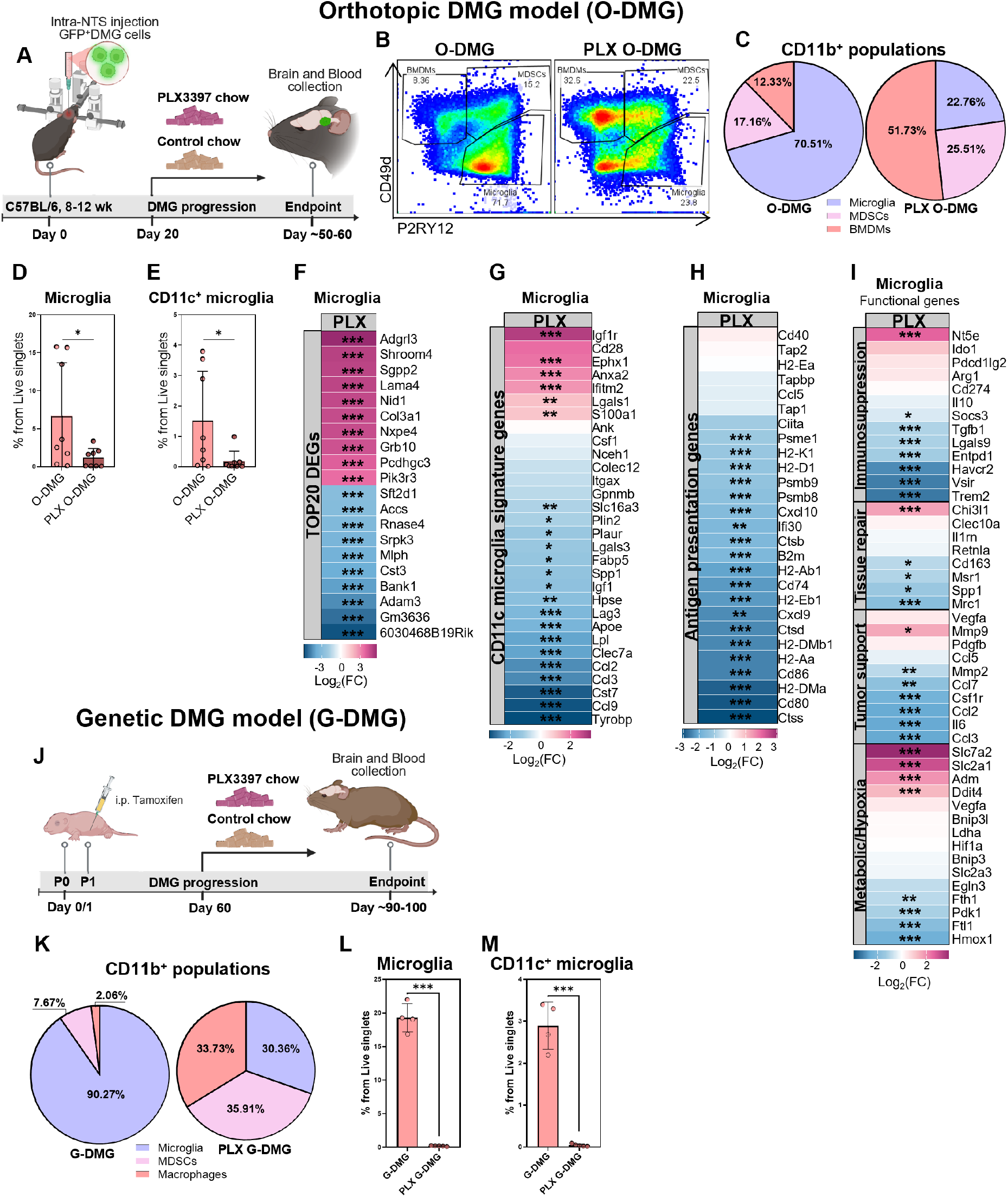
PLX3397 reprograms the myeloid compartment in DMG tumours, partially depleting microglial compartment, particularly reducing immune suppression and antigen presentation by decreasing CD11c^+^ microglial activation state. **A)** PLX3397 treatment scheme in orthotopic (O-DMG) murine model. **B**) Flow cytometry analysis performed to assess microglia in healthy HB/MB and its corresponding O-DMG tumour region: gating strategy of macrophages, myeloid derived suppressor cells (MDSCs) and microglia; **C**) Venn diagram showing percentages of microglia, macrophages and MDSCs from CD11b^+^ myeloid cells; **D**) the percentage of microglia from live singlets; **E**) the percentage of CD11c^+^ microglia from live singlets; n=4-8/group. Bulk RNA-sequencing of microglia (CD11b^+^CD49d^-^P2RY12^+^) sorted from DMG tumour region and its corresponding healthy hindbrain/midbrain (HB/MB): **F**) top 20 differentially expressed genes (DEGs); **G**) Functional immunosuppression, tissue repair, tumour support and metabolic/hypoxia associated gene heatmap; **H**) CD11c^+^ microglial gene signature heatmap; **I**) Antigen presentation gene signature heatmap; n=4/group. **J**) PLX3397 (PLX) treatment regime in the genetic (G-DMG) murine model. Flow cytometry analysis performed to assess microglia in untreated versus PLX-treated G-DMG whole brains: **K**) Venn diagram showing percentages of microglia, macrophages and MDSCs from CD11b^+^ myeloid cells; **L**) the percentage of microglia from live singlets; **M**) Percentages of CD11c^+^ microglia from live singlets; n=4-5/group. Significance: *p<0.05; **p<0.005; ***p<0.001, otherwise non-significant.

Preclinical studies have shown that CSF1R blockade can modulate myeloid populations and enhance T-cell activity in adult glioma^55,58^. Pexidartinib is clinically approved for tenosynovial giant cell tumour and achieves effective target inhibition *in vivo*^59–62^. Moreover, Pexidartinib/PLX3397 is currently in clinical trials for use alone or in combination with radiation therapy and temozolomide for the treatment of recurrent glioblastoma patients^63,64^. We therefore used PLX3397 (further referred to as PLX) to assess how CSF1R inhibition perturbs myeloid–T cell interactions in DMG. Similar to above, we used the immunocompetent orthotopic and genetic DMG models to assess the effect of the CSF1R inhibitor. O-DMG mice were given the diet containing PLX 20 days post-tumour implantation (**Fig. 5A**) and at day 60 post-tamoxifen induction in G-DMG (**Fig. 5J**). In both models PLX was therefore administrated at a time point when DMG tumours are starting to establish.

PLX markedly remodelled the brain myeloid compartment (**Fig. 5B-C; Supplementary 10**). Overall proportion of CD11b⁺ myeloid cells was significantly depleted in the O-DMG tumour after the PLX treatment (**Supplementary 10A**). After the PLX treatment, the myeloid compartment composition shifted: O-DMG-associated microglia were strongly reduced (from 70.51% to 22.76%) and no longer dominated the CD11b⁺ myeloid compartment (**Fig. 5C-D**). Meanwhile, the relative proportions of BMDMs and MDSCs increased within the myeloid compartment compared with microglia, despite unchanged infiltration into O-DMG tumours, indicating a change in the composition of the myeloid compartment rather than increased recruitment (**Fig. 5C, Supplementary 10B-C**). Notably, which was confirmed by flow cytometry showing significantly reduced CD11c⁺ microglia subpopulation in O-DMG tumours after the PLX treatment (**Fig. 5E**). Bulk RNA sequencing of O-DMG-associated global microglia population (**Fig. 5F, Supplementary 11A**) revealed a decreased CD11c^+^ microglia signature after the PLX treatment (**Fig. 5G**). Moreover, after the PLX treatment, O-DMG-associated microglia exhibited decreased antigen presentation (**Fig. 5H**) and immunosuppressive gene signatures in the remaining microglial population (**Fig. 5I**). Enrichment analyses including TTRUST transcription factor network, Gene Ontology Molecular Function and Reactome pathway analyses further indicated that after partial depletion, residual O-DMG-associated microglia adopt a less immune-engaged state, with suppression of core immune regulators, including *Spi1, Nfkb1* and *Irf8* (**Supplementary 11B-C**), alongside enrichment of extracellular matrix organization, cytoskeletal and adhesion-related pathways, with concurrent downregulation of antigen processing and presentation, rRNA processing and translation programs (**Supplementary 11D-G**). Collectively these findings suggest reduced immune-cell crosstalk and a partial reversion towards a quiescent, tissue-maintaining microglial phenotype.

To further assess the impact of PLX on the myeloid compartment remodelling, we have validated several of these key findings in the G-DMG model (**Fig. 5J**). Similarly to the O-DMG model, the overall proportion of CD11b⁺ myeloid cells was significantly depleted in the G-DMG tumour after the PLX treatment (**Supplementary 10D**). G-DMG tumour-associated microglia decreased from approximately 90% to 30% within the CD11b⁺ myeloid compartment after the PLX treatment (**Fig. 5K-M**). Moreover, similarly to the O-DMG model, in G-DMG tumours, CD11c⁺ microglia subset was significantly decreased (**Fig. 5M).** Together, these data show that CSF1R inhibition perturbed the myeloid compartment, with a partial reduction in the global microglial population and diminished CD11c^+^ microglia gene signatures. Consistent with this, PLX treatment also altered systemic cytokine profiles (**Supplementary 12**), including increased CX3CL1 and IL-13 (**Supplementary 12S, 12H**) and trends toward reduced CXCL9 and CCL2 (**Supplementary 12U, 12N**). These changes suggest that CSF1R inhibition reshapes chemokine networks governing myeloid cell recruitment, reduces antigen presentation and immunosuppressive signalling within the TME, suggesting that DMG-associated microglia may have a reduced capacity to sustain chronic T cell activation.

### CSF1R inhibition partially rescues DMG-associated T cell exhaustion-like dysfunctional state

Given that CSF1R inhibition with PLX perturbs the myeloid compartment, resulting in partial depletion of global microglia population, including a decrease in antigen presentation and immune suppression signatures as well as a reduction in CD11c^+^ microglia, we next sought to determine whether these changes were associated with altered T cell states. To further assess the impact of PLX-mediated myeloid perturbation on T-cell exhaustion, we have phenotyped the T cells in G-DMG and O-DMG models.

Untreated G-DMG tumours exhibited low CD4/CD8 ratio indicating relative CD8^+^ T cell dominance compared to CD4^+^ T cells (**Supplementary 2B**). PLX3397 treatment shifted the CD4/CD8 ratio toward a more balanced immune profile by increasing the relative abundance of CD4+ T cells while retaining CD8+ T-cell dominance (**Supplementary 13A**). Despite these changes, the total abundance of either of G-DMG-associated T cell subsets did not significantly change after the PLX treatment (**Supplementary 13B-D**). However, the phenotyping of G-DMG tumour-associated CD8^+^ T cells (**Fig. 6A**) revealed increased repeated antigen experience associated activation molecule CD44 after PLX, suggesting increased T cell activation. Interestingly, G-DMG tumour-associated CD8^+^ T cells displayed increased expression of inhibitory PD-1 (**Fig. 6A**), which could be recognized as a preventive switch in the cytotoxic T cells from chronic stimulation and terminal exhaustion. Whereas, CD4^+^ T cells have displayed increased expression of inhibitory TIM-3, decreased PD-1 and activation related CD44 after the PLX treatment (**Fig. 6B**). Considering CD4^+^ T cell-mediated activation of CD8^+^ T cells, these findings suggest partial rejuvenation of CD4^+^ T cells, which goes in hand with observed increased activation and partially relieved exhaustion of CD8^+^ T cells. Collectively, these findings suggest a partial restoration of T cell functionality and alleviation of exhaustion associated states after PLX-induced remodelling of the myeloid compartment.

**Fig. 6.**
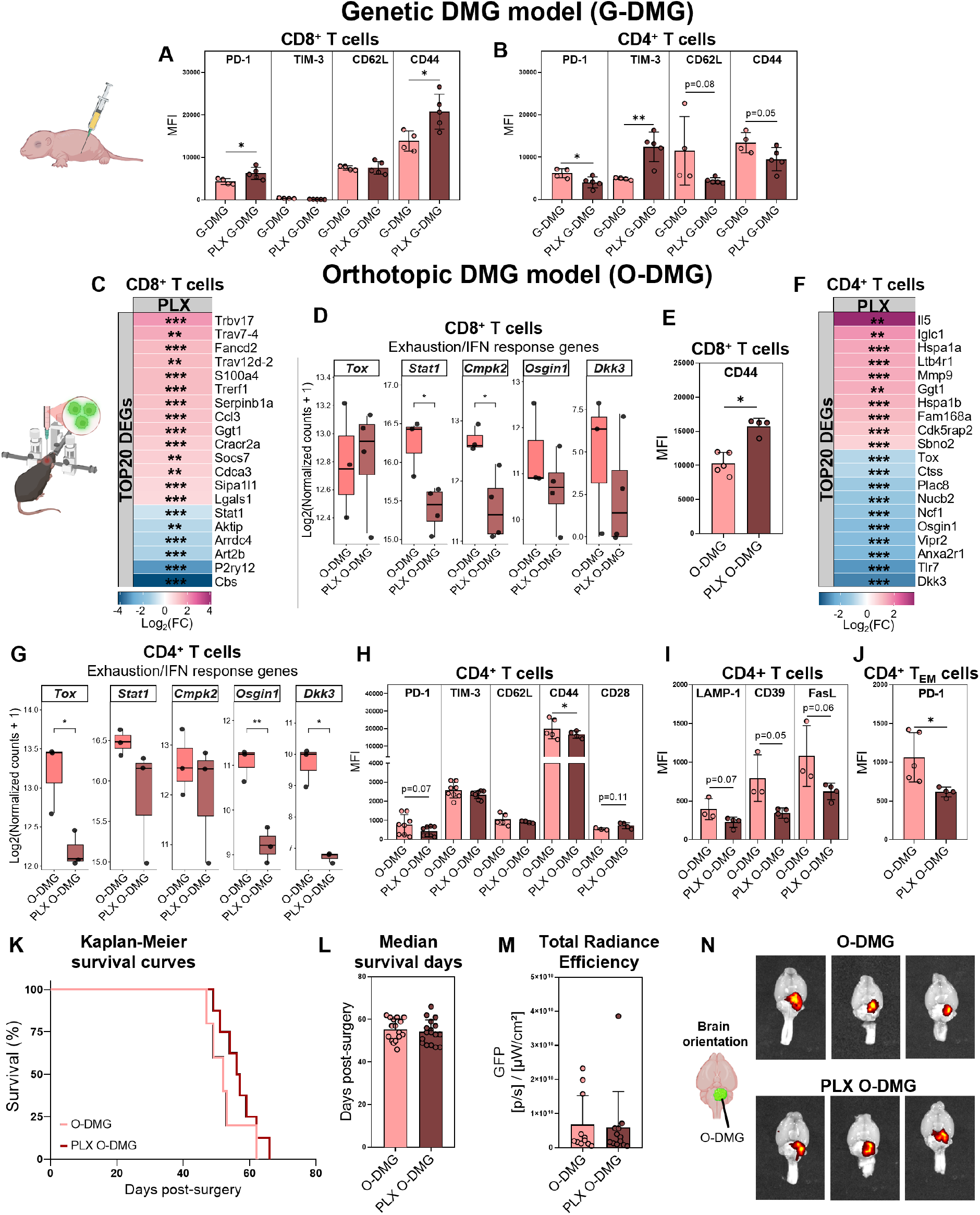
Changes in DMG-associated T cells after PLX3397 treatment alleviated T cell exhaustion. Flow cytometry analysis performed to assess T cells in untreated versus PLX3397-treated G-DMG (PLX G-DMG) whole brains. Median fluorescence intensity (MFI) of PD-1, TIM-3, CD62L, CD44 in: **A**) CD8^+^ T cells and **B**) CD4^+^ T cells. Bulk RNA-sequencing of CD8^+^ T cells sorted from untreated versus PLX3397-treated O-DMG tumour region: **C**) top 20 differentially expressed genes (DEGs); **D**) exhaustion and interferon response-related genes; n=3-4/group. **E**) Flow cytometry analysis of CD44 protein levels on CD8^+^ T cells, expressed as MFI. Bulk RNA-sequencing of CD4^+^ T cells sorted from untreated versus PLX3397-treated O-DMG tumour region: **F**) top 20 differentially expressed genes (DEGs); **G**) exhaustion and interferon response-related genes; n=3/group. Flow cytometry analysis performed to assess CD4^+^ T cells in untreated and PLX3397-treated O-DMG tumours: **H**) median fluorescence intensity (MFI) of PD-1, TIM-3, CD62L, CD44 and CD28; **I**) MFI of surface LAMP-1, CD39 and FasL; **J**) MFI of PD-1 from CD4^+^ T_EM_ cells. Comparisons between untreated and PLX3397-treated O-DMG tumour-bearing mice: **K**) Kaplan-Meier survival curves; **L**) Median survival days, n=16-18/group. **M**) Total radiance efficiency of GFP signal from DMG tumour measured with Caliper Life Sciences IVIS Lumina II, n=11-12/group. **N**) Representative images of GFP signal measurements (visualized as spectral overlay mask) in untreated and PLX3397-treated O-DMG brains. Significance: *p<0.05; **p<0.005; ***p<0.001, otherwise non-significant.

Next, we validated these key findings in the O-DMG model. Contrary to the G-DMG model, in the O-DMG, the tumour-associated T cell compartment was dominated by CD4^+^ T cells, resulting in a high CD4/CD8 T cell ratio (**Supplementary 2B**). PLX3397 treatment shifted this ratio towards a more balanced T cell distribution by increasing the relative proportion of O-DMG tumour-associated CD8^+^ T cells, while preserving CD4^+^ T cell predominance (**Supplementary 13E**), without significantly affecting the overall abundance of either T cell subset (**Supplementary 13F-H**). Moreover, PLX treatment did not significantly alter the distribution of CD4^+^ or CD8^+^ T cell naïve, central and effector-memory subsets in the O-DMG tumours (**Supplementary 13I-J**).

O-DMG-associated CD8^+^ T cells exhibited a transcriptionally and functionally reprogrammed state after PLX treatment (**Fig. 6C; Supplementary 14A).** At the transcriptional level, PLX treatment was associated with selective CD8^+^ T cell clonal expansion, reflected by increased T cell receptor (TCR) variable region transcripts (*Trbv17, Trav7-4, Trav12d-2*; **Fig. 6C**), alongside upregulation of TCR signalling and survival-associated genes, such as *Cracr2a, Kcna3, Blrc6, Serpinb1a, Ccnd3*, supporting enhanced activation and persistence (**Supplementary 14B-C**). Moreover, after the PLX treatment, O-DMG tumour-associated CD8^+^ T cells have displayed reduced chronic interferon response genes (*Stat1, Cmpk2*; **Fig. 6D**). TTRUST (**Supplementary 14D-E**), GO Molecular Function (**Supplementary 14F-G**) and Reactome enrichment analyses (**Supplementary 14H-I**) indicated that after PLX treatment, O-DMG-associated CD8^+^ T cells adopt an activated, stress-adaptive effector state, characterized by enrichment of Heat Shock Transcription Factor 1 (HSF1)-associated and E26 transformation-specific (ETS) family regulatory networks, cytokine signalling, and programmed cell death pathways, alongside suppression of innate immune, G protein-coupled receptor (GPCR), and endosomal/TLR-associated signalling programs. On the protein level, O-DMG tumour-associated CD8^+^ T cells showed increased repeated antigen experience associated activation molecule CD44 in PLX-treated DMG tumours, suggesting increased T cell activation (**Fig. 6E)**, despite no changes in the expression of other inhibitory, activation, cytotoxicity or exhaustion-related molecules such as PD-1, CD39, granzyme B, FasL and LAMP-1 (**Supplementary 15**).

Transcriptomic analysis of the global CD4⁺ T cell population in untreated and PLX-treated O-DMG tumours (**Fig. 6F; Supplementary 16A**) revealed that CD4⁺ T cells exhibited enhanced stress response, activation and migratory programs alongside suppression of exhaustion and dysfunction-associated signalling, including reduced *Tox* and *Irf8*, with increased *Stat6* and *Fosl2* activity (**Fig. 6G; Supplementary 16B**). Complementarily, TTRUST transcription factor network (**Supplementary 16D-E**), Gene Ontology (**Supplementary 16F-G**), and Reactome analyses (**Supplementary 16H-I**) further supported this transition, revealing enrichment of stress-response, kinase signalling, cytokine signalling, and migratory/remodelling pathways, together with suppression of innate immune response, IFN response, antigen presentation, endosomal, and TLR-associated programs. Moreover, at the protein level, the global CD4⁺ T cell population (**Fig. 6H-J**), predominantly the effector (CD4⁺ T_EM_) cells, showed a significant reduction in PD-1 expression after PLX treatment (**Fig. 6J, Supplementary 17A**). CD4⁺ T cells also exhibited a trend towards decreasing surface LAMP-1, CD39 and FasL protein expression levels, suggesting reduced exhaustion-associated features following PLX-mediated myeloid perturbation (**Fig. 6I, Supplementary 17B-C**), suggesting partial alleviation of the exhaustion and inhibitory signalling in the helper T cells.

The reproducibility and uniformity of the O-DMG model enabled us to determine whether PLX-induced myeloid remodelling and the associated partial rejuvenation of T cells translated into prolonged survival or reduced tumour burden. PLX diet did not significantly affect the survival of O-DMG tumour-bearing mice (**Fig. 6K-L**) or the tumour burden indicated by the GFP total radiance efficiency (**Fig. 6M-N; Supplementary 17D**). However, there was a significant decrease in the spleen weights observed in PLX-treated mice compared to untreated O-DMG-bearing or healthy mice at the end of the experiment (**Supplementary 17E**) with no major differences in the overall body weight measurements over time between the untreated and PLX-treated O-DMG-bearing mice groups (**Supplementary 17F**). Although partial microglial depletion and remodelling of the myeloid compartment did not significantly reduce tumour burden or improve survival, these findings indicate that this approach alone is insufficient for tumour control. However, the partial reversal of T cell exhaustion, including reduced PD-1 expression, suggests that microglia contribute to T cell dysfunction in DMG. Rather than broadly depleting microglia, selectively targeting microglia–T cell interactions may therefore represent a more effective therapeutic strategy and could enhance the efficacy of immune checkpoint blockade.

Together, these findings indicate that PLX-induced changes in the tumour-associated myeloid compartment, to some extent alleviated exhaustion/dysfunction in DMG-associated CD8^+^ and to greater extent in CD4^+^ T cells, promoting more activated and stress-adaptive states (**Fig. 7**). While these changes were insufficient to significantly affect tumour burden or animal survival as a standalone treatment, they suggested that reversal of microglia-driven T cell suppression and dysfunction may enhance T cell functionality within the DMG TME, supporting an adjunctive strategy aimed at overcoming key barriers limiting effective DMG treatment.

**Fig. 7.**
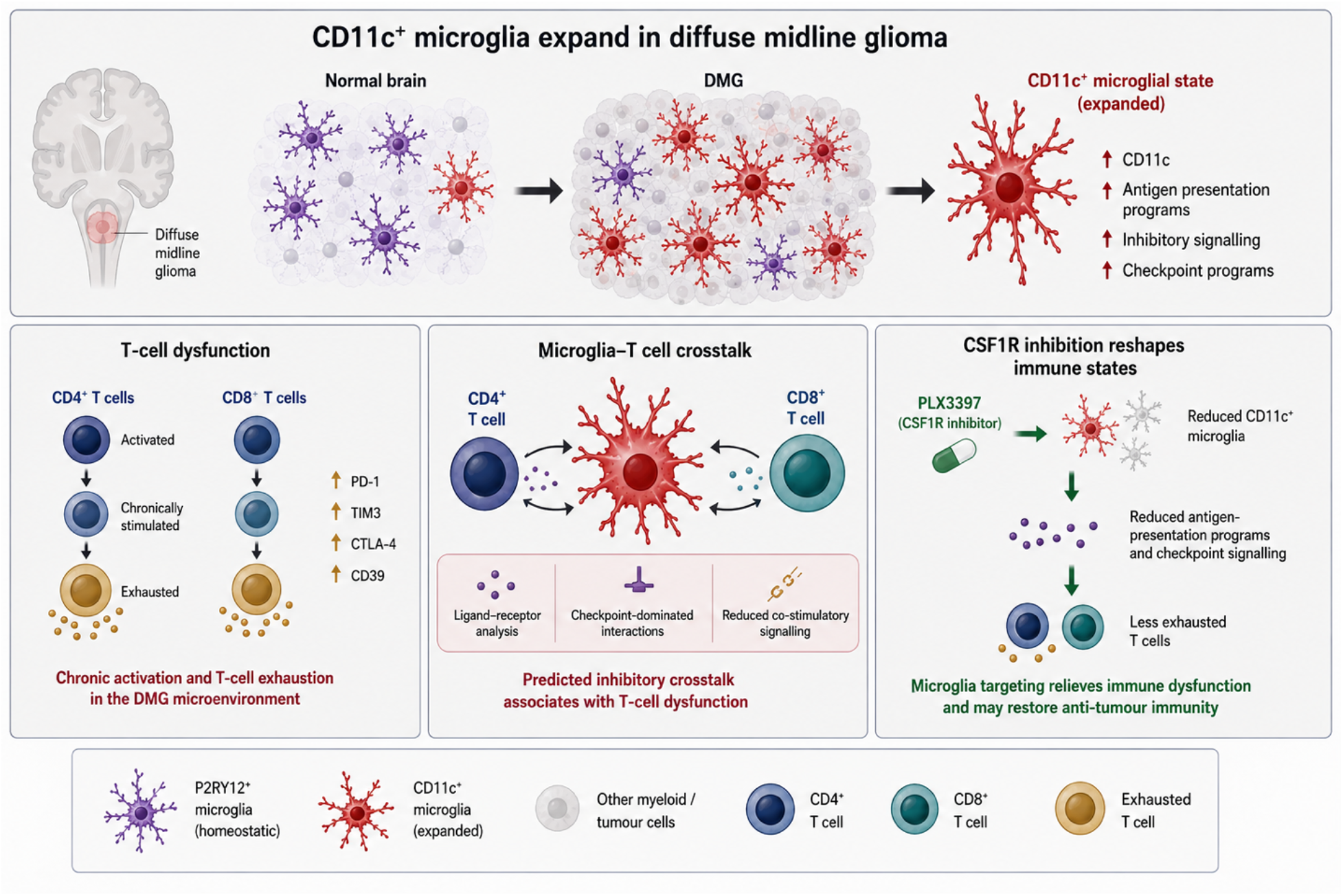
CD11c^+^ microglia are associated with T cell dysfunction in diffuse midline glioma (DMG). DMG promotes the expansion of a CD11c⁺ microglial activation state characterized by increased antigen presentation-associated, inhibitory, and checkpoint-related transcriptional programs. DMG-infiltrating CD4⁺ and CD8⁺ T cells exhibit features of chronic stimulation and exhaustion, including increased expression of PD-1, TIM-3, CTLA-4, and CD39. Ligand– receptor analyses identified predicted inhibitory crosstalk between microglia and T cells together with reduced co-stimulatory signalling, suggesting that CD11c⁺ microglia contribute to immune dysfunction within the DMG microenvironment. Pharmacological inhibition of CSF1R with PLX3397 depleted global microglia and reduced CD11c⁺ microglial state, decreased antigen presentation-associated and checkpoint-related programs, which was associated with a shift toward a less exhausted T-cell phenotype. Together, these findings identify a microglia–T cell axis that contributes to immune dysfunction in DMG and may represent a target for immunomodulatory therapies. This image was generated with AI assistance using ChatGPT (version 5) under human supervision and review.

## Discussion

Our results reveal a novel microglia–T cell axis in DMG, characterised by antigen presentation in the absence of canonical co-stimulation and associated with T cell dysfunction. Resolving tumour-associated myeloid populations by lineage, we identify microglia rather than infiltrating macrophages as the dominant population linked to this phenotype and further demonstrate that a CD11c⁺ microglial state is enriched for inhibitory signalling and primed to contribute to dysfunctional T cell engagement.

### Microglia and macrophages adopt distinct activation states in DMG

Microglia and bone marrow–derived macrophages (BMDMs) are ontogenetically distinct myeloid populations within the glioma microenvironment^32^. Many previous reports have broadly referred to tumour-associated myeloid cells (TAMs) without resolving these distinct populations. Microglia are long-lived, self-renewing residents of the CNS derived from yolk sac progenitors, whereas BMDMs arise from bone marrow hematopoietic stem cells and are recruited to tumours via the blood^65–70^. Although both populations adopt similar phenotypes in response to tumour-derived cues, they retain lineage-imprinted differences: microglia maintain homeostatic CNS-associated signatures, while BMDMs exhibit enhanced migratory, inflammatory, and antigen-presentation programs^28,32,71^.

We applied P2RY12 and CD49d to resolve TAM heterogeneity in diffuse midline glioma (DMG). Previous work in adult glioblastoma, demonstrated that these populations can be confidently distinguished using lineage-specific markers: P2RY12 is robustly expressed on resident microglia, whereas CD49d (ITGA4) identifies infiltrating BMDMs. Importantly, these markers were validated against lineage-traced populations, confirming that they reflect ontogeny rather than activation state^28,32^. This approach allowed us to accurately discriminate resident microglia from infiltrating BMDMs. Previous studies in DMG have established that TAMs represent major component of the TME and largely exhibit immunosuppressive and tumour-supportive phenotypes^8,30^. However, these studies primarily considered TAMs as a single population without fully resolving lineage specific differences.

Our observations indicate that both microglia and BMDMs acquire antigen-presenting, activated, and immunosuppressive states within the tumour microenvironment. These findings expand our prior understanding of the tumour microenvironment in DMG^72–74^ and glioblastoma^28^, further supporting the necessity of tumour-derived cues to drive functional convergence while preserving lineage-specific programs. Importantly, our data extend these observations by showing that microglia and macrophages do not simply converge into a single TAM state but instead retain distinct activation states despite overlapping functional features. Taken together with the existing DMG literature, these findings highlight that resolving ontogeny reveals an additional layer of heterogeneity that is not captured when TAMs are analysed as a unified population.

### CD11c^+^ microglia activation state is associated with inhibitory signalling

CD11c (ITGAX) in microglia is best interpreted as an activation-associated state marker rather than a fixed lineage-defining feature. Across development and disease, CD11c⁺ microglia emerge transiently in contexts of increased cellular turnover, inflammation, or tissue remodelling, and are characterized by enhanced phagocytic activity, antigen presentation capacity, and immunomodulatory functions^41–45^. In developmental systems, CD11c expression tracks with apoptotic burden and can revert when local cues subside, supporting its dynamic and reversible nature^52^. Similarly, in neuroinflammatory and neurodegenerative settings such as Experimental Autoimmune Encephalomyelitis (EAE), disease-associated CD11c⁺ microglia state align with broader activation programs involving phagocytosis, lipid metabolism, and immune regulation, including *Itgax, Apoe, Lpl,* and *Clec7a*^75,76^, and have been implicated in antigen presentation and T cell–modulating functions in inflammatory CNS environments^42,44,45^.

Here, we identify and characterize previously unrecognized disease-associated CD11c^+^ microglia activation state emerging within the DMG TME. We show that CD11c^+^ microglial activation state is enriched for antigen presentation machinery and immunoregulatory signalling programs, including inhibitory pathways such as PD-L1, suggesting a direct capacity to influence local T cell behaviour^42,77–79^. This study reveals directly assess potential cellular crosstalk, ligand–receptor interaction mapping between CD4⁺ and CD8⁺ T cells and myeloid populations revealed multiple predicted interaction axes involving microglia, supporting the possibility of direct microglia–T cell communication within the TME. Our findings therefore refine the current understanding of myeloid heterogeneity in DMG and implicate this population as a potential contributor to local immune suppression.

Functionally, our perturbation data provide direct support for a role of CD11c⁺ microglia in shaping T cell states. Partial depletion of microglia using PLX decreases the abundance of CD11c⁺ microglia within live singlets and reduces the CD11c⁺ microglial signature among global microglia alongside broader antigen presentation and immunosuppressive programs. Strikingly, this is accompanied by a reduction in CD4⁺ and CD8⁺ T cell exhaustion signatures, indicating that microglia contribute to the establishment or maintenance of T cell dysfunction in DMG. Together, these effects suggest that the CD11c⁺ microglial state is potentially a key mediator of immunosuppressive microglia–T cell interactions, but this would require further validation.

### Implications for immunotherapy resistance in DMG

Our study provides a detailed characterization of the TME in unresectable paediatric DMG, highlighting features that may underlie the limited efficacy of GD2 or B7-H3 targeting CAR-T cell therapies currently applied in clinical trials^8,9^. Consistent with prior reports, we observe that DMG tumours are shaped by a highly immunosuppressive microenvironment, dominated by activated microglia, increased macrophage infiltration, and T cells exhibiting markers of chronic activation and exhaustion.

Notably, our analysis reveals that both resident and infiltrating immune cells display suppressive phenotypes. T cells within the tumour express gene signatures and surface proteins indicative of chronic stimulation and functional exhaustion yet retain intrinsic cytotoxic potential. This paradox suggests that T cells are capable of executing effector functions but are not effectively engaged in tumour cell killing, potentially due to inadequate activation signals or impaired antigen presentation.

We observed that in DMG tumours there is a high CD4/CD8 ratio, reflecting a relative paucity of CD8⁺ T cells compared to CD4⁺ T cells. This imbalance suggests a weakened anti-tumour immune response, with fewer cytotoxic effectors available to eliminate tumour cells, consistent with the observed exhaustion of both CD4⁺ and CD8⁺ T cell populations. Furthermore, this is in line with the immunosuppressive phenotype identified within the CD4⁺ T cell compartment, which may further contribute to impaired anti-tumour immunity^11–13,80^.

Our findings implicate macrophages and microglia as key mediators of immunosuppression in DMG. Despite evidence of antigen presentation within the TME, T cells appear unable to mount effective anti-tumour responses. We hypothesize that microglia contribute to this dysfunction by engaging T cells in “suppressory” synapses, prolonging interactions that fail to provide activating cues. This is supported by our observation that canonical co-stimulatory pathways, including CD40–CD40L and CD28–CD80/CD86, are not effectively engaged, potentially preventing full T cell activation.

A major strength of this study is the ability to reproduce our findings across two independent immunocompetent models of DMG. This is particularly important in the context of DMG research, where many preclinical studies rely on xenograft models implanted into immunodeficient hosts such as NOD-SCID mice. While these models are valuable for studying tumour biology, the absence of a functional immune system limits their ability to faithfully model tumour–immune interactions. By using immunocompetent models, we were able to interrogate endogenous myeloid and T cell responses within the native TME, providing a more physiologically relevant framework for understanding immune dysfunction in DMG.

Together, these results suggest that the DMG TME actively restrains T cell functionality through multiple complementary mechanisms, including suppressive antigen-presenting cells and disrupted co-stimulatory signalling. Such insights may explain why CAR-T therapies, despite trafficking to the tumour and recognizing antigens, exhibit limited clinical efficacy. Targeting the immunosuppressive microenvironment—particularly microglial and macrophage-mediated suppression—could therefore represent a critical strategy to enhance T cell-mediated immunotherapy in DMG. Although these changes were insufficient to significantly affect tumour burden or animal survival as a monotherapy, they suggested that reversal of microglia-driven T cell suppression and dysfunction may represent a potential therapeutic avenue and serve as a pre-requisite for developing effective DMG treatment strategies. Although these changes were insufficient to significantly affect tumour burden or animal survival as a monotherapy, they suggested that reversal of microglia-driven T cell suppression and dysfunction may represent a potential therapeutic avenue and serve as a pre-requisite for developing effective DMG treatment strategies. Future studies should further investigate whether therapeutic strategies that reverse microglia-mediated T-cell dysfunction effectively and restore T-cell activation and cytotoxicity, either alone or in combination with other immunotherapies, can translate into meaningful anti-tumour efficacy and survival benefit in DMG.

### Limitations

Several limitations of this study should be acknowledged. First, our conclusions are based primarily on transcriptional and phenotypic analyses, and direct functional validation of microglia–T cell interactions, for example through co-culture or antigen-specific assays, will be required to establish causality. Second, the use of bulk RNA sequencing limits resolution of cellular heterogeneity within defined populations. Third, CSF1R inhibition has systemic effects that may influence immune responses beyond the TME. Finally, validation of these findings in human DMG samples will be essential to confirm their translational relevance.

### Conclusions

In summary, our study identifies a previously undiscovered role for microglia in shaping T cell dysfunction in DMG and highlights the importance of resolving myeloid ontogeny and activation states within the tumour microenvironment. By implicating a CD11c⁺ microglial activation state in this process, we redefine the role of microglia in DMG and provide an urgently needed new framework for understanding how immune responses are constrained in DMG. This implicates microglia–T cell interactions as promising targets to enhance immunotherapeutic strategies in this otherwise intractable disease.

## Methods

### Animal Husbandry and Ethical Approval

Female mice were housed under specific pathogen-free conditions on a 12 h light/dark cycle with controlled temperature (21 ± 1°C) and humidity (55–60%), with ad libitum access to standard rodent chow and water. All animal procedures were approved by the Animal Experimentation Ethics Committee (AEEC) at University College Cork and the Health Products Regulatory Authority (HPRA) under project authorization number AE19130/P217 and were conducted in accordance with European Directive 2010/63/EU and S.I. No. 543 of 2012.

#### Orthotopic Mouse Model of DMG (O-DMG)

Mouse DMG cells were generated by intrauterine electroporation of *H3f3a^K27M^*, *Pdgfra^D842V^*, and DN*p53* constructs into the developing brainstem of C57BL/6 embryos at embryonic day 13.5, as previously described^48,49^. Tumours were harvested, dissociated into single-cell suspensions, and maintained ex vivo prior to implantation.

#### Cell Culture and Preparation

Mouse DMG cells were cultured under 5% CO_2_ at 37°C in tumour stem medium (TSM) consisting of DMEM/F12 and Neurobasal medium supplemented with B27 without vitamin A (all from Invitrogen), recombinant human bFGF (20 ng/mL), EGF (20 ng/mL), PDGF-AA (10 ng/mL), and PDGF-BB (10 ng/mL) (all from Shenandoah Biotechnology), and heparin (2 μg/mL; STEMCELL Technologies). Cell cultures were routinely screened for mycoplasma contamination using the Venor™GeM Mycoplasma Detection Kit (Minerva Biolabs). Immediately prior to implantation, cells were prepared as a single-cell suspension in sterile DPBS.

#### Tumour Implantation

Female C57BL/6 mice (model code: 057; Inotiv, UK) of 8-12 weeks old were anaesthetized with isoflurane (5% induction and 1.7–2.2% maintenance) and administered pre-operative subcutaneous injection of meloxicam (5mg/kg). Animals were positioned in a stereotaxic frame and, under aseptic conditions, 5 × 10^5^ DMG cells in a total volume of 5 μL were injected into the pontine region using stereotactic coordinates relative to lambda (AP −0.8 mm, ML −1.0 mm, DV −4.7 mm). Cells were infused at a rate of 2 μL/min. The needle was allowed to remain in place for 2 min prior to injection and for at least 2.5 min following injection before being slowly withdrawn to minimize reflux.

Following surgery, the incision was closed with sutures and mice were recovered on a warming pad. Post-operative analgesia (meloxicam, 1 mg/kg) was provided in the drinking water for 4 days, and animals were monitored daily. DMG tumours are lethal within approximately 50-60 days post-implantation. Mice were euthanized upon reaching predefined humane endpoints, including moderate neurological/motor symptoms, ≥20% body weight loss or signs of significant distress.

#### Genetic Model of DMG (G-DMG)

A tamoxifen-inducible *Nestin-CreER^T2^* genetically engineered mouse model of DMG (G-DMG) was used as previously described^2^. This model drives expression of H3F3A-K27M and mutant PDGFRA together with conditional deletion of *Tp53* in Nestin-expressing neural progenitor cells following tamoxifen-induced Cre recombination.

Tamoxifen (Sigma, T5648) was dissolved in corn oil (Sigma, C8267) at a concentration of 5 mg/mL by incubation at 37°C. The solution was sterilized through a 0.22 μm filter, protected from light, and stored at 4°C for up to 7 days. Cre recombinase activity was induced by intraperitoneal administration of tamoxifen (3 mg/40 g body weight) using a 30-gauge insulin syringe. Male and female neonatal mice received tamoxifen on postnatal days 0 and 1, with injections administered 24 h apart. Mice were euthanized at predefined humane endpoints, as DMG tumours lead to mortality approximately 90–100 days after tamoxifen induction.

#### PLX3397 Treatment

Pexidartinib (PLX3397; Boc Bioscience, B0084-470807) was incorporated into AIN76A chow diet (Ssniff) at 600 ppm. Control animals received matched AIN76A chow without PLX3397. Mice were maintained on control chow upon arrival and allowed to acclimatize for one week prior to tumour implantation. Twenty days following orthotopic implantation, mice were either maintained on control chow or switched to PLX3397-containing chow for maximum of 40 days post-surgery.

### Survival and body weight analysis

Survival was defined as the time from surgery to the onset of predefined humane endpoints associated with O-DMG progression, characterised by tumour growth and the development of moderate behavioural symptoms as assessed by established monitoring criteria. Survival curves were generated using Kaplan–Meier survival analysis. Differences between groups were analysed using the Log-rank test. Median survival times were calculated for each group. Animal body weights were recorded prior to surgery and every second day till endpoint. At each timepoint, differences in body weight between groups were assessed by Mann-Whiney U test. To account for multiple comparisons across timepoints, p-values were adjusted using Holm-Šidák method.

### Blood and tissue collection

Mice were sacrificed by cervical dislocation between 9am and 12pm. Blood was immediately collected from the heart in heparinized tubes (368495, BD) and centrifuged for 10 minutes at 1600 rpm at 4°C after multiple inversion of the tubes. Supernatant was collected and centrifuged again for 3 minutes at 13200 rpm at 4°C. Plasma was collected and stored at –80°C for further analysis. The pellet with blood cells and collected lymph nodes were immediately used for flow cytometry analysis. Spleen was collected, weighed and immediately used for flow cytometry analysis. Brain was collected and used for tumour size measurement via *In Vivo* Imaging System (IVIS) (see section “Tumour size measurement via *In Vivo* Imaging System (IVIS Lumina Series II)”) prior to further processing via flow cytometry analysis.

### Tumour size measurement via *In Vivo* Imaging System (IVIS Lumina Series II)

At the humane endpoint (HE), mice were sacrificed and brains were carefully collected for DMG tumour visualization with IVIS Lumina Series II equipment and Living Image software (Caliper/Perkin Elmer, Akron, OH, USA). *Ex vivo* brains were initially imaged from the ventral surface, then bisected along the sagittal plane into left and right hemispheres and re-imaged with the cut surfaces facing upward to expose the internal tumour. GFP-expressing DMG tumours were visualized by epifluorescence imaging using the GFP filter set (445–490 nm excitation, 515–575 nm emission). Composite images were generated using the image overlay function on Living Image. The brain tumour region was defined around the brainstem along the midline. Regions of interest (ROIs) were drawn over this area and quantified as Total Radiant Efficiency (TRE) in the calibrated unit Radiant Efficiency [p/s/sr]/[µW/cm^2^], which is normalized by the software for differences in exposure time. Tumour TRE was calculated by subtracting the background fluorescence measured in the same ROI. The data was obtained and analysed using Living Image version 4.5.2 and 4.7.2 software (https://www.perkinelmer.com/uk/lab-products-and-services/resources/in-vivo-imaging-software-downloads.html).

### Perfusion and microscopy of GFP^+^ O-DMG tumours

Female mice were anesthetized with intraperitoneal injection of Pentobarbital Sodium (Abbeyville Vet Hospital, 90mg/kg of body weight) and subjected to transcardiac perfusion with PBS, followed by 4% w/v paraformaldehyde (PFA) (Fisher Scientific, Cat. P/0840/53). Brain tissue was extracted and post-fixed in PFA overnight at 4°C, and then transferred to 25% w/v sucrose (S/8600/60, Fisher Scientific) for a minimum of 48h at 4°C. Following complete saturation, brains were snap frozen in isopentane (2-Methylbutane, VWR Chemicals BDH, Cat. 294524E) cooled with liquid nitrogen and stored at -80°C until processing. The brains were cryosectioned into 30µm tissue sections with a cryostat (Leica Biosystems, California, USA). The tissue sections were air-dried and mounted in ProLong Diamond Antifade Mountant (Thermo Fisher Scientific, Cat. P36984), visualized with VS200 slide scanner (Evident Scientific, Tokyo, Japan).

### Flow cytometry analysis

Mice were sacrificed by cervical dislocation, brain was harvested in ice-cold HBSS, spleen and lymph nodes were harvested in ice-cold DPBS, blood was collected into heparinized tubes. All samples were kept on ice until processing.

The brain was surgically divided into non-tumour (NTR) and tumour (DMG) regions, with appropriate healthy brain tissue– forebrain (FB) and midbrain/hindbrain (MB/HB). The tissues were transferred to C tubes and mixed with enzymes from Mouse tumour dissociation kit (Miltenyi Biotec, Cat. 130-096-730) in DMEM media and processed using m_impTumor_02 program on the gentleMACS Octo Dissociator (Miltenyi Biotec, Cat. 130-096-427). Then the tissues were enzymatically digested for 40 minutes at 37°C, afterwards processing with m_impTumour_03 program on the dissociator. Dissociated cell suspension was filtered via 40-μm cell strainers (Fisher Scientific, Cat. 11587522), centrifuged and subjected to percoll gradient. Obtained cell pellets was washed and resuspended in 0.5% BSA and 1 mM EDTA in DPDB.

Blood cells were obtained after plasma isolation (described in “Blood and tissue collection”) and subdued to blood cell lysis with RBC lysis buffer (Invitrogen, Cat. 00-4300-54) for 10 minutes at room temperature, washed with DPBS and pelleted, further resuspended in 0.5% BSA and 1 mM EDTA in DPDB.

Cervical lymph nodes were dissociated using 40-μm cell strainers, washed with ice-cold DPBS and pelleted, further resuspended in 0.5% BSA and 1 mM EDTA in DPDB.

All samples were further incubated with TruStain FcX™ PLUS (anti-mouse CD16/32) (Biolegend, Cat. 156604) and Fixable Live/Dead Alexa Fluor 488 viability die (Invitrogen, Fisher Scientific, Cat. L34970) to prevent nonspecific binding and discriminate dead cells, respectively. Then samples were stained with surface antibody cocktails (**Table 1**). These stainings were washed after staining and fixed in 2% of methanol-free paraformaldehyde in DPBS (made from 16% stock solution, Thermo Scientific Pierce, Cat. 28906) for 30min at RT, washed, and resuspended in FACS buffer. Samples for intracellular staining of Ki-67, granzyme B, and perforin were fixed and permeabilized using the Foxp3/Transcription Factor Staining Buffer Set (Invitrogen eBioscience, Thermo Fisher Scientific; Cat. No. 50-112-8857) according to the manufacturer’s instructions. Following staining, all samples were acquired by flow cytometry within 24 hours of preparation.

**Table 1.** Antibody used in the flow cytometry analysis with BD FACSCelesta and FACSSymphony analysers.

| Antigen | Conjugate | Clone | Manufacturer | Cat. number |
| --- | --- | --- | --- | --- |
| B220/CD45R | APC-Cy7 | RA3-6B2 | BioLegend | #103224 |
| CD107a/LAMP-1 | Brilliant Violet 605 | 1D4B | BioLegend | #121643 |
| CD11b | Brilliant Violet 750 | M1/70 | BioLegend | #101267 |
| CD11b | Alexa Fluor 488 | M1/70 | BioLegend | #101217 |
| CD11c/ITGAX | PE | N418 | BioLegend | #117308 |
| CD19 | Alexa Fluor 700 | 1D3/CD19 | BioLegend | #152414 |
| CD28 | Brilliant Violet 421 | E18 | BioLegend | #122023 |
| CD39 | PE-Dazzle59 | Duha59 | BioLegend | #143812 |
| CD4 | APC | RM4-5 | BioLegend | #100516 |
| CD40 | APC-Cy7 | 3/23 | BioLegend | #124638 |
| CD40 | Brilliant Violet 605 | 3/23 | BD | #745218 |
| CD40L/CD154 | PE | SA047C3 | BioLegend | #157004 |
| CD44 | Brilliant Violet 605 | IM7 | BioLegend | #103047 |
| CD45 | PerCP | 30-F11 | BioLegend | #103130 |
| CD49d | PE-Dazzle594 | R1-2 | BioLegend | #103626 |
| CD62L | Pacific Blue | W18021D | BioLegend | #161208 |
| CD8A | Brilliant Violet 785 | 53-6.7 | BioLegend | #100750 |
| CD86/B7-2 | PerCP | GL-1 | BioLegend | #105026 |
| CLEC7A/CD369 | Brilliant Violet 605 | 218820 | BD | #749793 |
| CTLA-4/CD152 | Brilliant Violet 605 | UC10-4B9 | BioLegend | #106323 |
| FasL/CD178 | PE | MFL3 | BioLegend | #106606 |
| Granzyme B | Alexa Fluor 700 | QA16A02 | BioLegend | #372222 |
| PD-1/CD279 | PE-Dazzle594 | RMP1-30 | BioLegend | #109116 |
| PD-L1/CD274 | Brilliant Violet 786 | 10F.9G2 | BioLegend | #124331 |
| CD4 | Brilliant Violet 605 | RM4-5 | BioLegend | #100545 |
| CD4 | APC | RM4-5 | BioLegend | #100516 |
| TIM-3 | Alexa Fluor 700 | 215008R | R&D systems | #FAB1529RN |
| TLR2/CD282 | Brilliant Violet 421 | CB225 | BioLegend | #148605 |
| TLR2/CD282 | PE | 6C2 | Invitrogen | #12-9021-82 |
| TREM2 | APC | 237920 | R&D systems | #FAB17291A |
| Ki-67 | APC-Cy7 | 11F6 | BioLegend | #151232 |
| KLRG1/MAFA | APC-Cy7 | 2F1/KLRG1 | BioLegend | #138426 |
| Ly6C | PerCP | HK1.4 | BioLegend | #128028 |
| Ly6C | Pacific Blue | HK1.4 | BioLegend | #128014 |
| Ly6G | Brilliant Violet 605 | 1A8 | BioLegend | #127639 |
| MHC-II/ I-A/I-E | Alexa Fluor 700 | M5/114.15.2 | BioLegend | #107622 |
| P2RY12 | APC | S16007D | BioLegend | #848006 |
| P2RY12 | APC-Cy7 | S16007D | BioLegend | #848024 |
| Perforin | Brilliant Violet 421 | S16009A | BioLegend | #154319 |
| ST2 | APC-Cy7 | PK136 | BD | #746876 |

Samples were acquired on a BD FACSCelesta (BD Biosciences) using V5, B4, and R3 detection channels. Obtained data was analysed with FlowJo (version 10.0.8). Dead cells and doublets were gated out prior to downstream analysis. Cellular phenotypes were assessed using high parameter flow cytometry panels (**Fig. S1**) containing markers to identify cell types and activation states. For the estimation of median fluorescence intensity (MFI), minimum 2000 events were used.

### Sorting of immune populations for bulk RNA sequencing

C57BL/6 mouse brains were harvested and dissected as described in section ‘Flow cytometry analysis’ and further digested in gentleMACS C tubes (Miltenyi Biotec) and dissociated by using enzymes in Tumour dissociation kit with an automatic tissue gentleMACS Dissociator (Miltenyi Biotec), and subsequently incubating the samples for 40min at 37°C. Digested brain tissue was mechanically dissociated via a 70-μm cell strainer with ice-cold 1X HBSS. Pelleted samples were washed with FACS buffer and subsequently incubated with LIVE/DEAD Fixable Green Dead Cell Stain Kit Alexa Fluor 488 (ThermoFisher Scientific, Cat. L34970) and TruStain FcX™ PLUS (anti-mouse CD16/32) (Biolegend, Cat. 156604) for 15 min and stained with an antibody cocktail (**Table 1**), including CD45, CD4, CD8, CD11b, P2RY12, CD49d (**Table 2**) and DAPI (Cat.40043, VWR) for 20 min on ice, kept away from light. After washing, the sample was passed through a 40-μm cell strainer and sorted in a 4-way sort with a “High quality 4-way sort” setting. Four cell populations were obtained (Microglia, Macrophages, CD4^+^ T cells and CD8^+^ T cells), rapidly checked for purity (**Fig. S1**) and immediately centrifuged, the pellet was rapidly lysed in RLT buffer and snap frozen on dry ice to preserve RNA (further described in “Sample preparation for bulk RNA sequencing”).

**Table 2.** Antibody used in the flow cytometry sorting experiment with BD FACSAria Fusion sorter.

| Antigen | Conjugate | Clone | Manufacturer | Cat. number |
| --- | --- | --- | --- | --- |
| CD11b | Alexa Fluor 488 | M1/70 | BioLegend | #101217 |
| CD4 | APC | RM4-5 | BioLegend | #100516 |
| CD45 | PerCP | 30-F11 | BioLegend | #103130 |
| CD49d | PE-Dazzle594 | R1-2 | BioLegend | #103626 |
| CD8A | Brilliant Violet 785 | 53-6.7 | BioLegend | #100750 |
| P2RY12 | APC | S16007D | BioLegend | #848006 |

### Sample preparation for bulk RNA sequencing

Sorted cells were pelleted at 450×g for 6min at 4°C, and immediately lysed in RLT Plus buffer (QIAGEN). Samples were vortexed for 1 min, snap-frozen on dry ice and stored at -80°C. Total RNA was extracted using the RNeasy Plus Micro Kit (QIAGEN, Cat. 100545) according to the manufacturer’s instructions and eluted in RNase-free water. Extracted total RNA was stored at - 80°C until the samples were shipped to Novogene for the Plant and Animal Eukaryotic Strand Specific mRNA (WOBI) service.

### Library preparation and bulk RNA sequencing

Library preparation and sequencing were performed by Novogene (WOBI stranded mRNA sequencing service). RNA quality and concentration were assessed prior to library preparation. Poly(A)-selected mRNA was enriched from total RNA, followed by fragmentation and reverse transcription to generate first-strand cDNA. Second-strand cDNA synthesis was performed, followed by end repair, A-tailing, and ligation of Illumina P5 and P7 adapters. Libraries were PCR-amplified, size-selected, and purified using magnetic bead-based clean-up. Library quality and fragment size distribution were assessed using Qubit fluorometry, quantitative PCR, and Bioanalyzer analysis. Libraries were pooled in equimolar ratios and sequenced as 150 bp paired-end reads on an Illumina NovaSeq 6000 platform.

### Bulk RNA-sequencing data processing and differential expression analysis

Data processing and quality control were performed following best-practice principles from nf-core RNA-sequencing pipelines, using Python v3.13.13) within a Conda (v24.11.3) managed environment. Raw FASTQ files were first subjected to quality control using FastQC (v0.12.1), summarized by MultiQC (v1.33) to assess sequencing quality, GC content, and adapter contamination. Adapter trimming and quality filtering were performed using Trim Galore! (v0.6.10), which incorporates Cutadapt (version 4.9). Sequencing adapters (including platform-specific P5 and P7 adapters) were removed, low-quality bases (Phred score < Q20) were trimmed from read ends, and reads shorter than 20 bp after trimming were discarded to avoid multi-mapping of short reads. Post-trimming quality control was performed using FastQC to confirm read quality and adapter removal efficiency. Transcript abundance was quantified using Salmon (v1.10.3) in quasi-mapping (pseudoalignment) mode against the mouse reference transcriptome (Ensembl release 113, GRCm39). This approach enables fast, bias-aware estimation of transcript-level expression without requiring genome alignment. Transcript-level quantifications were imported into R (v4.4.3) using the tximport package (v1.34.0) and summarized to gene-level counts based on Ensembl gene annotations. Gene-level count matrices were generated for all downstream analyses.

Differential gene expression analysis was performed using DESeq2 (v1.46.0) in R (v4.4.3). Given the exploratory nature of the study and limited sample size (n = 3–4 per group), results were prioritised based on nominal (unadjusted) p-values in combination with effect size (log_2_ fold change) to identify biologically relevant signals. Variance-stabilizing transformation (VST) was used for exploratory analyses, including principal component analysis. Log_2_ fold changes were shrinkage-adjusted using the apeglm method to improve interpretability of effect sizes. Key findings from the transcriptomic analysis were independently validated at the protein level using flow cytometry and multiplex cytokine assays, providing orthogonal confirmation of biologically relevant changes observed in the RNA-sequencing data.

### Cell-cell communication analysis

Gene identifiers were converted to *Mus musculus* Entrez Gene IDs using the Metascape annotation database (updated 2026-02-01), and duplicate mappings were collapsed prior to analysis. Overlap among the CD4^+^ T cells, CD8^+^ T cells, and Microglia gene sets was analysed in Metascape at both the direct gene level and the functional level based on shared enriched ontology terms. To reduce nonspecific associations, ontology terms containing more than 100 genes were excluded from the functional overlap analysis. Results were visualized using Circos plots generated by Metascape (metascape.org).

Predicted cell–cell communication was inferred using the CellChatDB.mouse ligand–receptor database. Differentially expressed ligands in sender cells were matched to cognate receptors in receiver cells, and interaction scores were calculated by combining ligand and receptor expression, differential regulation (log2 fold change), and statistical significance into a single interaction score, which was then aggregated for each sender–receiver pair to generate a cell–cell interaction matrix, which was visualized as a heatmap using pheatmap.

### Cytokine determination by LEGENDplex™

Cytokine levels in serum were determined using a custom LEGENDplex™ mouse panel focused on T cell activation/exhaustion13-plex panel, (Biolegend, Cat. No. 740446) and a predefined mouse Inflammatory panel (Biolegend, Cat. 740446); a bead-based assay that allows simultaneous measurement of analytes based on cell size using flow cytometry. Specifically, the cytokines measured were IL-33, CXCL9, CXCL10, IL-27, IL-23, IFN-γ, IFNIFN-β, IL-2, IL-6, IL-4, IL-10, GM-CSF, CX3CL1, IL-13, IL-1α, IL-1β, TGF-β1, IL-17a, CCL2, IL-12p70 and TNF-α. The assay was carried out in a 96-well plate following manufacturers’ instructions.

### Statistical analysis and data visualization

RNA-sequencing data analyses were performed in R (v4.4.3; RStudio) using DESeq2 (v1.46.0) for normalization and differential gene expression analysis. R session information and package versions are provided for reproducibility. Flow cytometry data, multiplex cytokine measurements (LEGENDplex^TM^), and *in vivo* experimental readouts (including body weight, spleen weight, and survival analysis) were analysed using GraphPad Prism (v8.0.1). Illustrations for the figures were created with Biorender.com. Data distribution was assessed using the Shapiro–Wilk test for normality. Depending on distribution, parametric data were analysed using unpaired Student’s t-tests and visualized as mean together with standard deviation (Mean ± SD), while non-parametric data were analysed using Mann–Whitney U tests, visualized as median (interquartilew range/IQR). Survival curves were analysed using the Mantel–Cox (log-rank) test. At each timepoint, differences in body weights longitudinal observations to account for multiple comparisons across timepoints, p-values were adjusted using Holm-Šidák method. A significance threshold of p<0.05 was applied throughout. Statistical significance was determined as defined in the statistical analysis section (+ p < 0.1; *p < 0.05; **p < 0.005; ***p < 0.001).

## Data availability

All scripts used for preprocessing, quantification, and downstream analysis are available in the Keane Lab GitHub repository, ensuring full reproducibility.

## Acknowledgements

We extend our appreciation to Colette Manley and Dr. Kenneth O’Riordan for their invaluable technical expertise and support. We also greatly appreciate Geomatrix, UAB (Lithuania) for providing access to computational infrastructure and server space for storage and analysis of high-dimensional bulk RNA-sequencing data.

## Authors information

LK and AB conceived the study. LK, AB and CCM designed experiments, interpreted the data and wrote the first draft of the manuscript. CCM, MMD, AB, JOR, GSST and BS performed DMG stereotactic surgeries. AB and CCM performed animal experiments, IVIS imaging, sample collection and processing for cytokine determination, flow cytometry, and bulk RNA sequencing. AB performed flow cytometry analysis and sorting of immune populations for bulk RNA sequencing and cytokine measurement using LEGENDplex in O-DMG. JH and AB performed flow cytometry analysis in G-DMG. MMH performed G-DMG experiments. AC, TP and EH generated orthotopic DMG animal model and provided DMG cells and expertise, SB and JDL provided the genetic DMG model. CF provided flow cytometry expertise and technical support, OOL, GMM, GC, JFC have critically evaluated the results and provided constructive feedback.

## Supplementary Figures

**Fig. S1:**
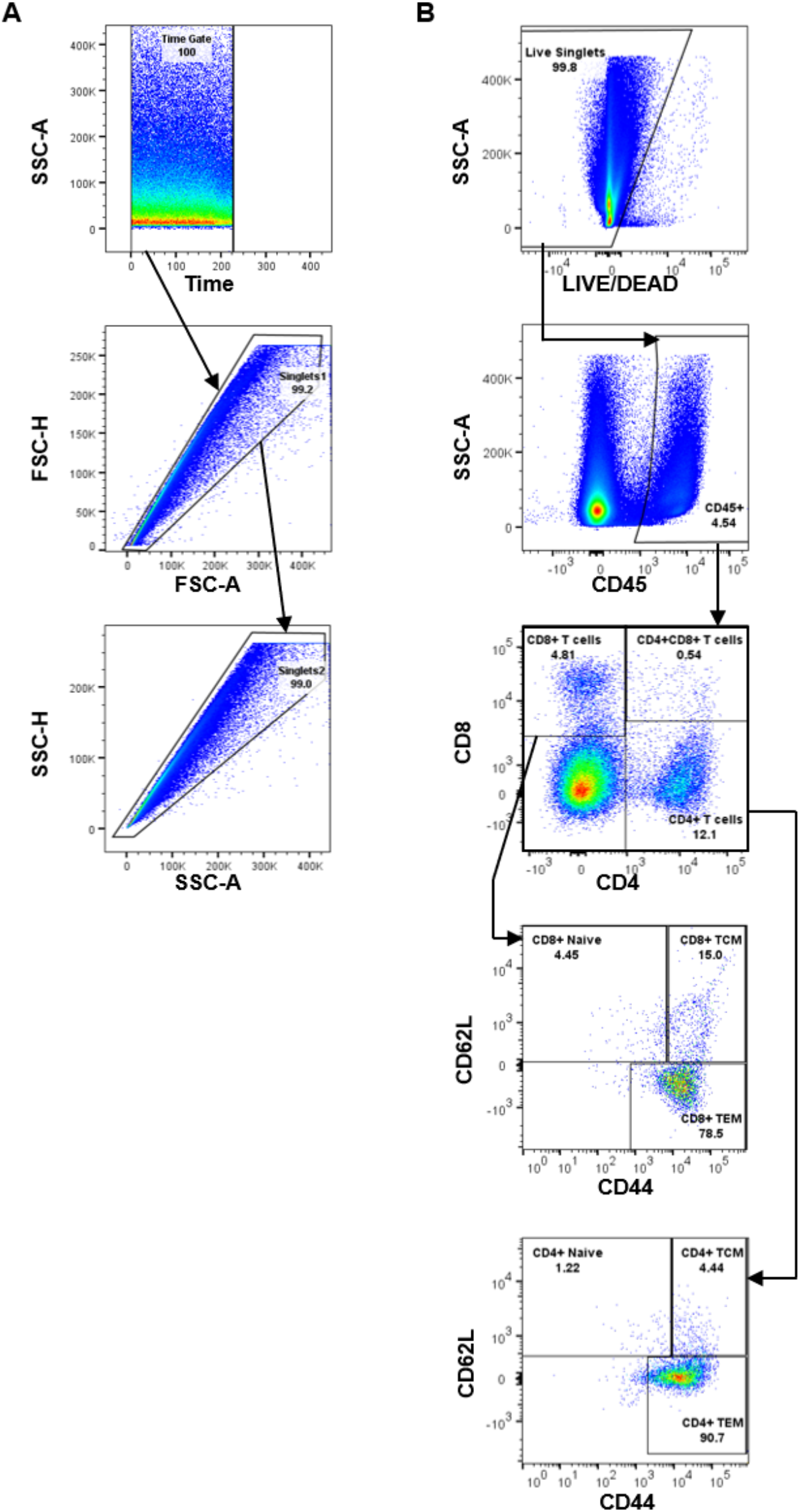
Flow cytometry gating strategy of brain immune populations. Gating strategy of immune populations was performed with FlowJo software: **A**) Gating strategy of single cells. Gated populations from live singlets: **B**) CD4^+^ and CD8^+^ T cell populations and naive, central-memory (T_CM_) and effector-memory (T_EM_) subsets.

**Fig. S2:**
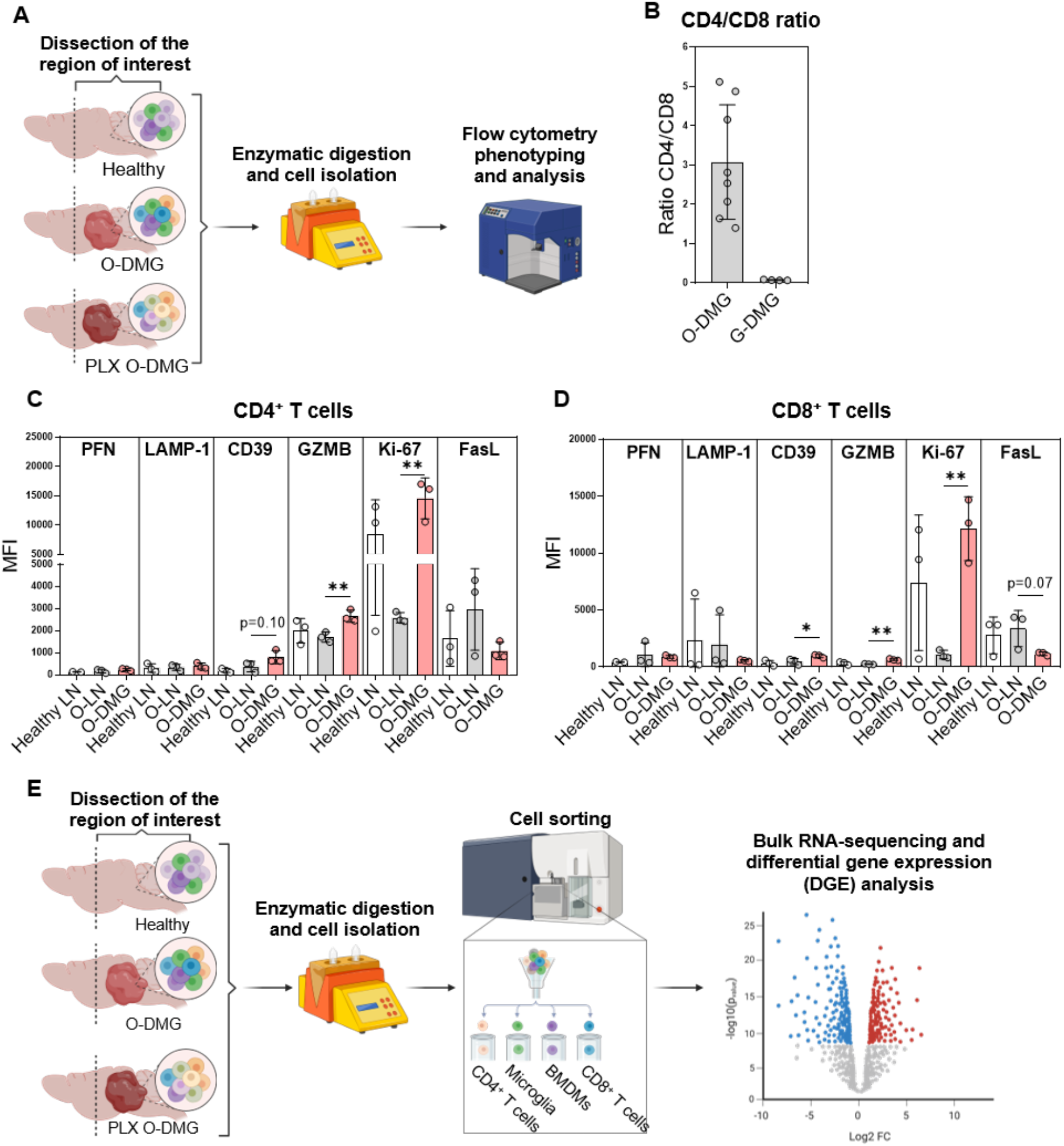
C57BL/6 experimental study design and T cell phenotyping in the orthotopic (O-DMG) and genetic (G-DMG) tumour models. **A**) Flow cytometry analysis and immune cell phenotyping in the regions of interest (healthy midbrain/hindbrain region versus O-DMG tumour region); **B**) CD4^+^ to CD8^+^ T cell ratio in G-DMG and O-DMG brains, n=4-8/group. O-DMG brain T cell expression of perforin (PFN), granzyme B (GZMB), Ki-67 and surface LAMP-1, CD39, FasL expressed as median fluorescence intensity (MFI) from: **C**) CD4^+^ T cells; **D**) CD8^+^ T cells. n=3/group *p<0.05; **p<0.005. **E**) Schematic of bulk RNA-sequencing of FACS-sorted immune populations: CD4^+^ and CD8^+^ T cells, microglia and macrophages sorted from healthy MB/HB, untreated and PLX3397-treated DMG tumours. Sorting was performed with BD FACSAria Fusion. Bulk RNA-sequencing was performed by Novogene.

**Fig. S3:**
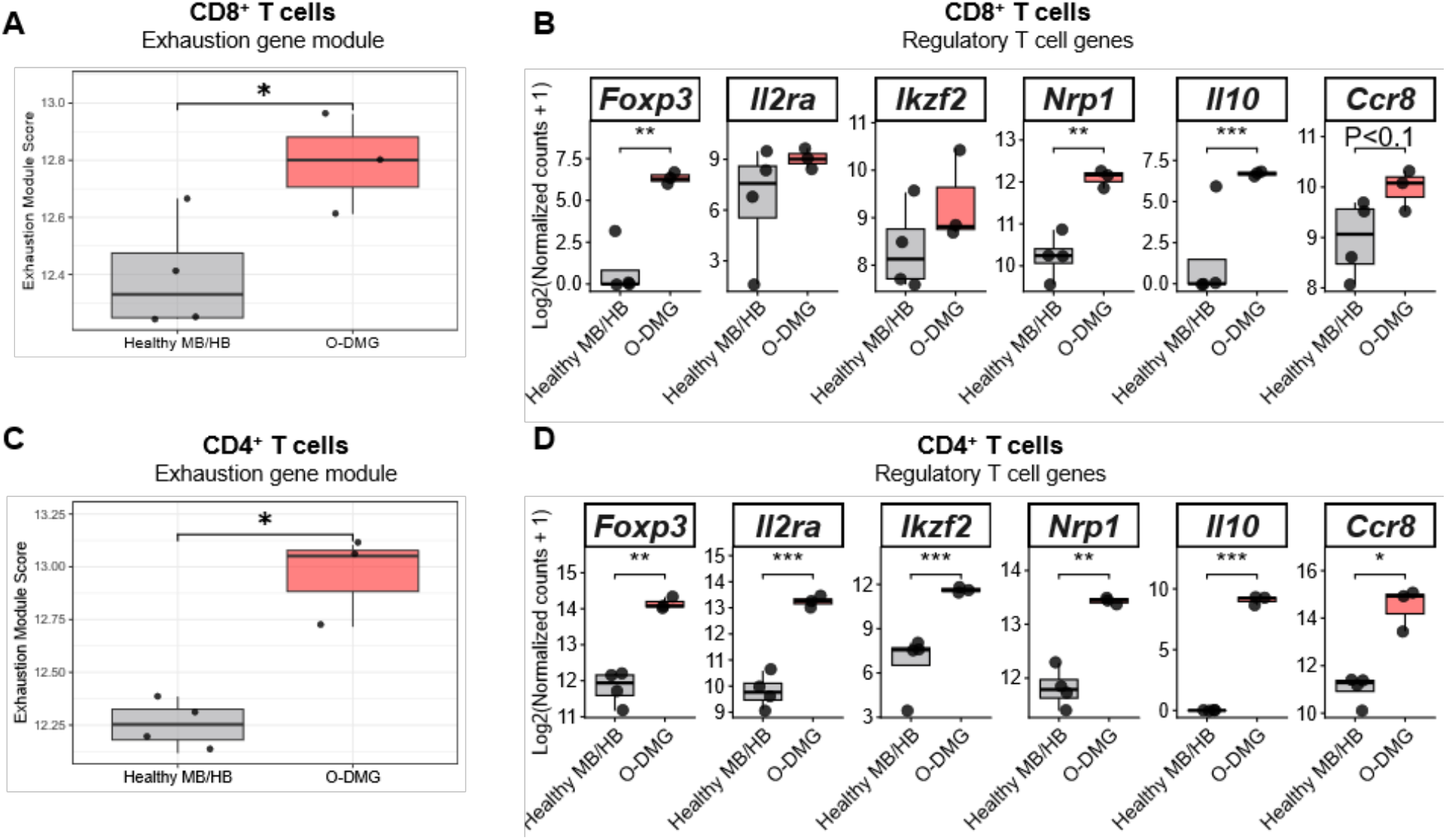
Transcriptomic phenotyping of T lymphocytes in the orthotopic DMG (O-DMG) tumours. CD4^+^ and CD8^+^ T cells sorted from healthy MB/HB, untreated O-DMG tumours. Sorting was performed with BD FACSAria Fusion. Bulk RNA-sequencing was performed by Novogene. **A**) Exhaustion gene cumulative module score in CD8^+^ T cells; **B**) Inhibitory/suppression-associated signalling genes in CD8^+^ T cells. **C**) Exhaustion gene cumulative module score in CD4^+^ T cells; **D**) Inhibitory/suppression-associated signalling genes in CD4^+^ T cells.

**Fig. S4:**
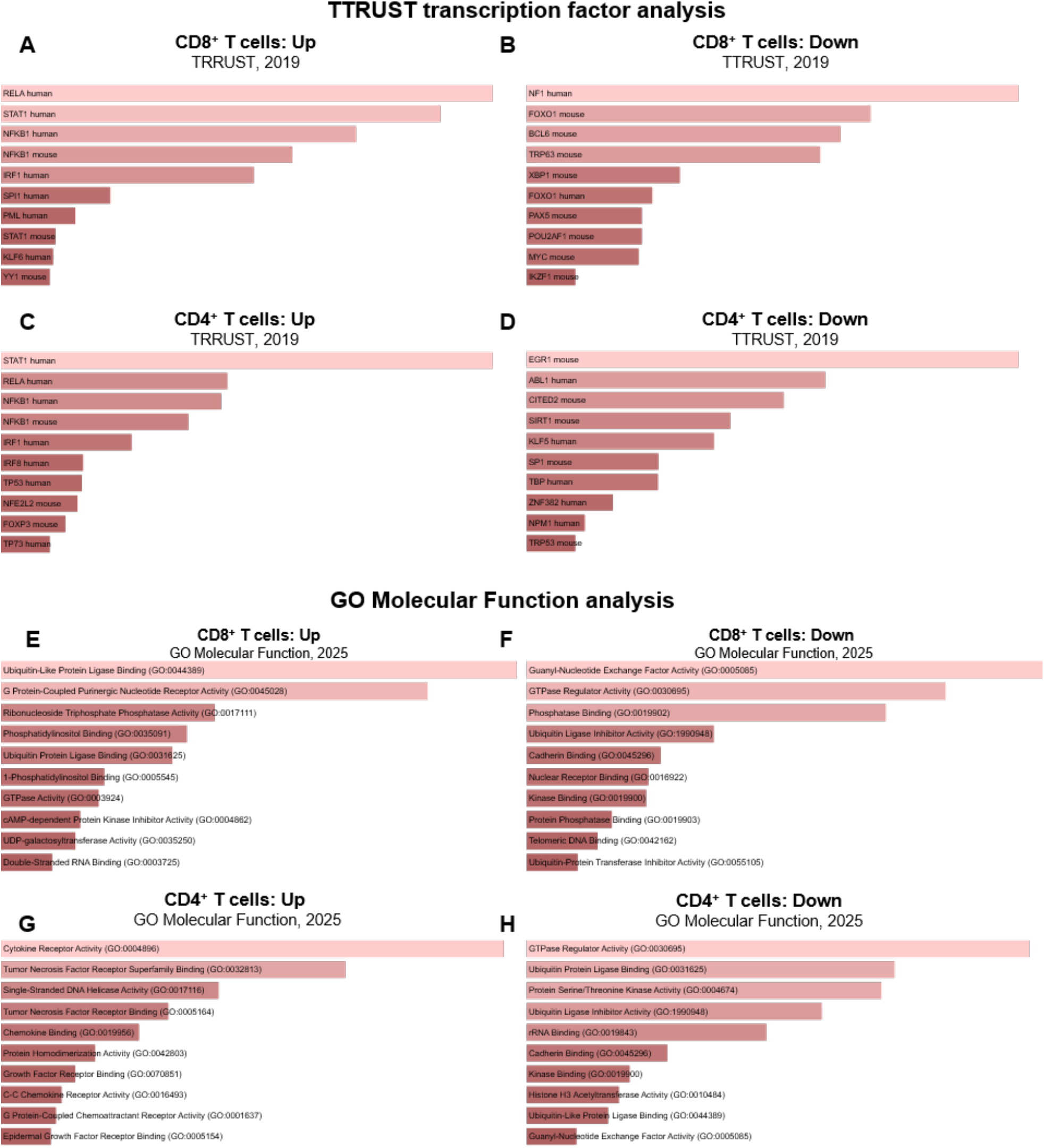
Enrichment analysis of sorted T lymphocytes from the orthotopic DMG (O-DMG) tumours. CD8^+^ and CD4^+^ T cells sorted from healthy MB/HB, untreated O-DMG tumours. Sorting was performed with BD FACSAria Fusion. Bulk RNA-sequencing was performed by Novogene. TTRUST transcription factor network analysis in T cells. CD8^+^ T cells: **A**) upregulated transcription factor networks; **B**) downregulated transcription factor networks. CD4^+^ T cells: **C**) upregulated transcription factor networks; **D**) downregulated transcription factor networks. GO Molecular Function pathway analysis in T cells. CD8^+^ T cells: **E**) upregulated; **F**) downregulated. CD4^+^ T cells: **G**) upregulated; **H)** downregulated. Only significant differentially expressed genes were used for these analyses, filtered with an unadjusted p<0.05 and log_2_ fold change >0.5.

**Fig. S5:**
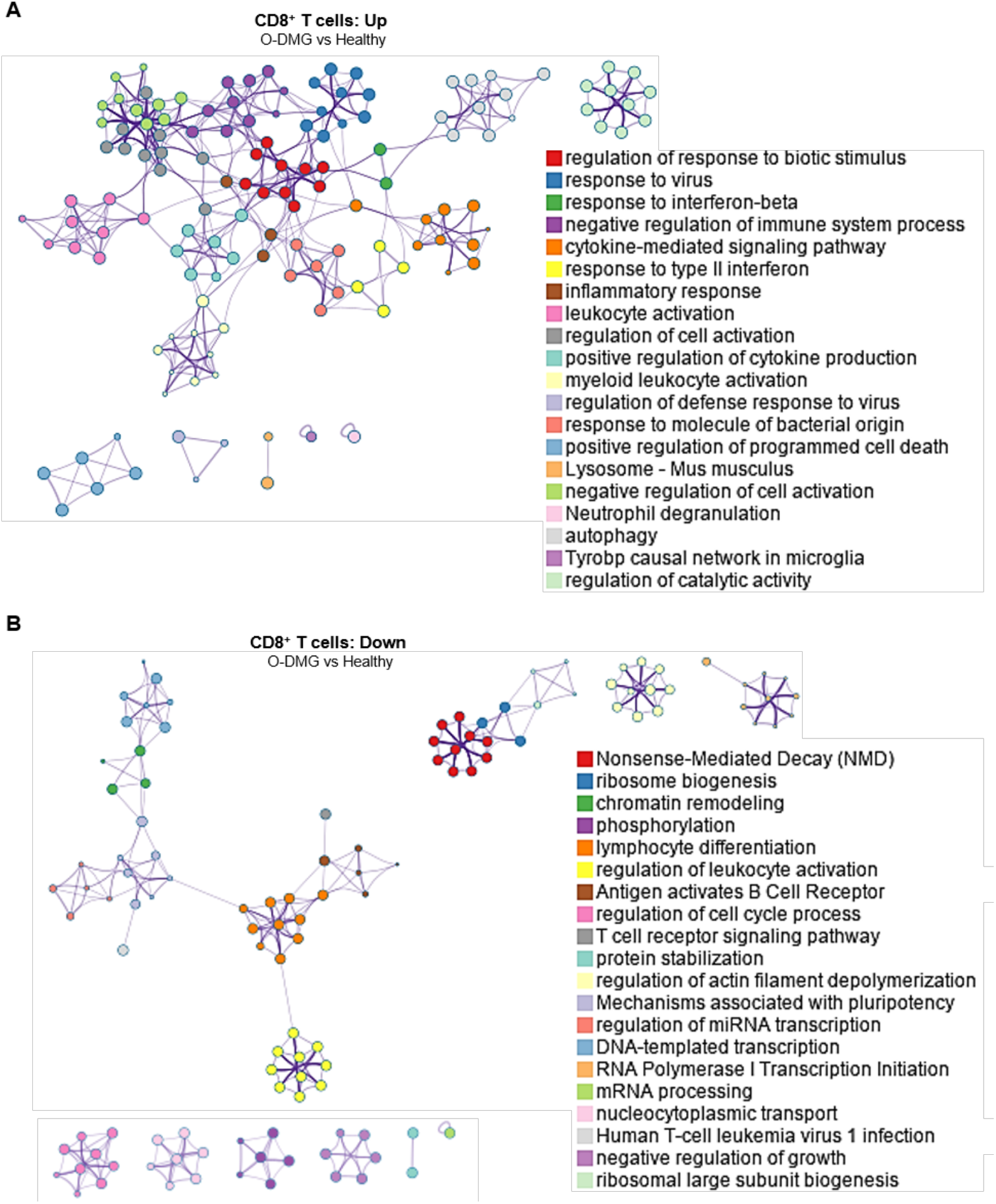
Biological processes enrichment analysis of sorted CD8^+^ T lymphocytes from the orthotopic DMG (O-DMG) tumours. CD8^+^ T cells sorted from healthy midbrain/hindbrain (MB/HB) regon and untreated O-DMG tumours. Sorting was performed with BD FACSAria Fusion. Bulk RNA-sequencing was performed by Novogene. The biological processes analysis wss performed with Metascape: **A**) upregulated and **B**) downregulated processed in O-DMG tumour-associated CD8^+^ T cells. Only significant differentially expressed genes were used for these analyses, filtered with an unadjusted p<0.05 and log_2_ fold change >0.5.

**Fig. S6:**
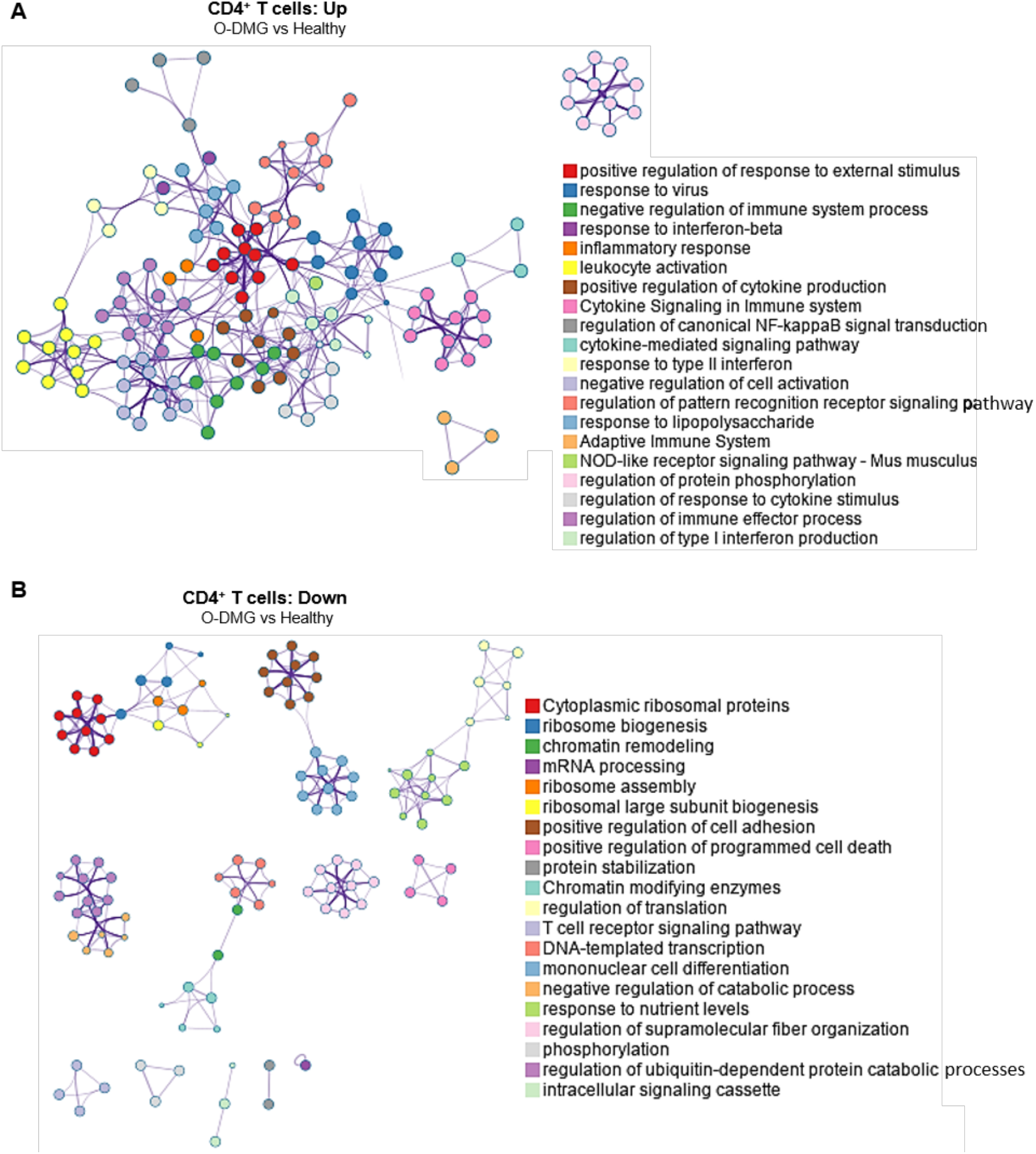
Biological processes enrichment analysis of sorted CD4^+^ T lymphocytes from the orthotopic DMG (O-DMG) tumours. CD4^+^ T cells sorted from healthy midbrain/hindbrain (MB/HB) regon and untreated O-DMG tumours. Sorting was performed with BD FACSAria Fusion. Bulk RNA-sequencing was performed by Novogene. The biological processes analysis wss performed with Metascape: **A**) upregulated and **B**) downregulated processed in O-DMG tumour-associated CD4^+^ T cells. Only significant differentially expressed genes were used for these analyses, filtered with an unadjusted p<0.05 and log_2_ fold change >0.5.

**Fig. S7:**
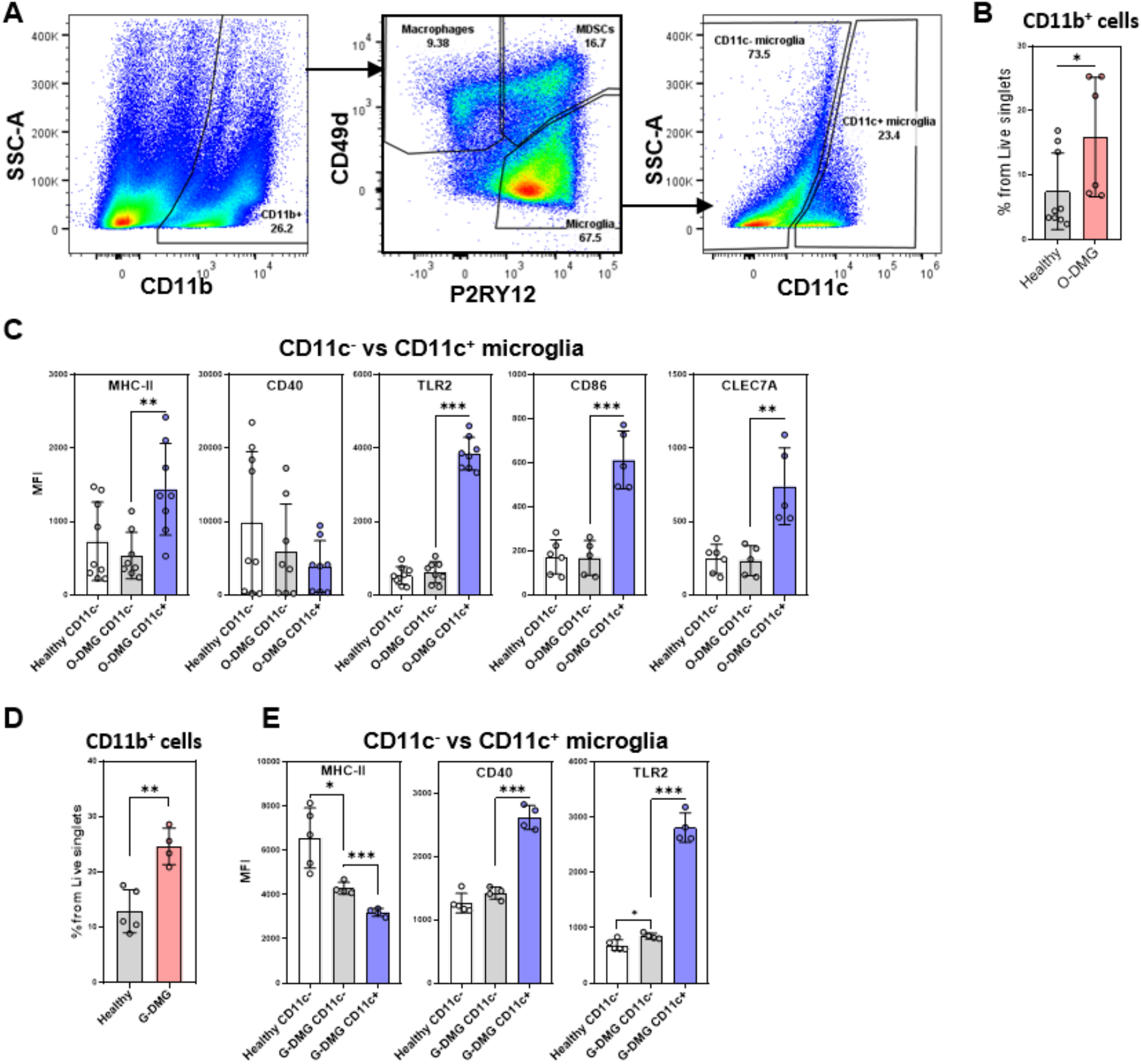
Flow cytometry phenotyping of myeloid immune populations in the orthotopic (O-DMG) and genetic (G-DMG) murine models. Flow cytometry phenotyping of myeloid populations in the healthy midbrain/hindbrain region versus O-DMG tumour region: **A**) Gating strategy of global CD11b^+^ myeloid cells, further gated into tumour-associated myeloid (TAMs) populations: macrophages (CD11b^+^CD49d^+^P2RY12^-^), microglia (CD11b^+^CD49d^-^ P2RY12^+^) and myeloid-derived suppressor cells (MDSCs; CD11b^+^CD49d^int^P2RY12^int^); global microglia population was further gated into CD11c^-^ and CD11c^+^ microglia. **B**) The percentage of CD11b+ myeloid cells taken from live singlets; **C**) Median fluorescence intensity (MFI) of MHC-II, CD40, TLR2, CD86, CLEC7A from healthy MB/HB and O-DMG tumour CD11c^-^ versus O-DMG-associated CD11c^+^ microglia. Flow cytometry of myeloid populations in the G-DMG tumours versus non-induced healthy whole brains: **D)** The percentage of CD11b+ myeloid cells taken from live singlets; **E**) MFI of MHC-II, CD40, TLR2 from healthy non-induced brain and genetic (G-DMG) tumour CD11c^-^ versus G-DMG-associated CD11c^+^ microglia; n=4-9/group. *p<0.05, **p<0.005, ***p<0.001.

**Fig. S8:**
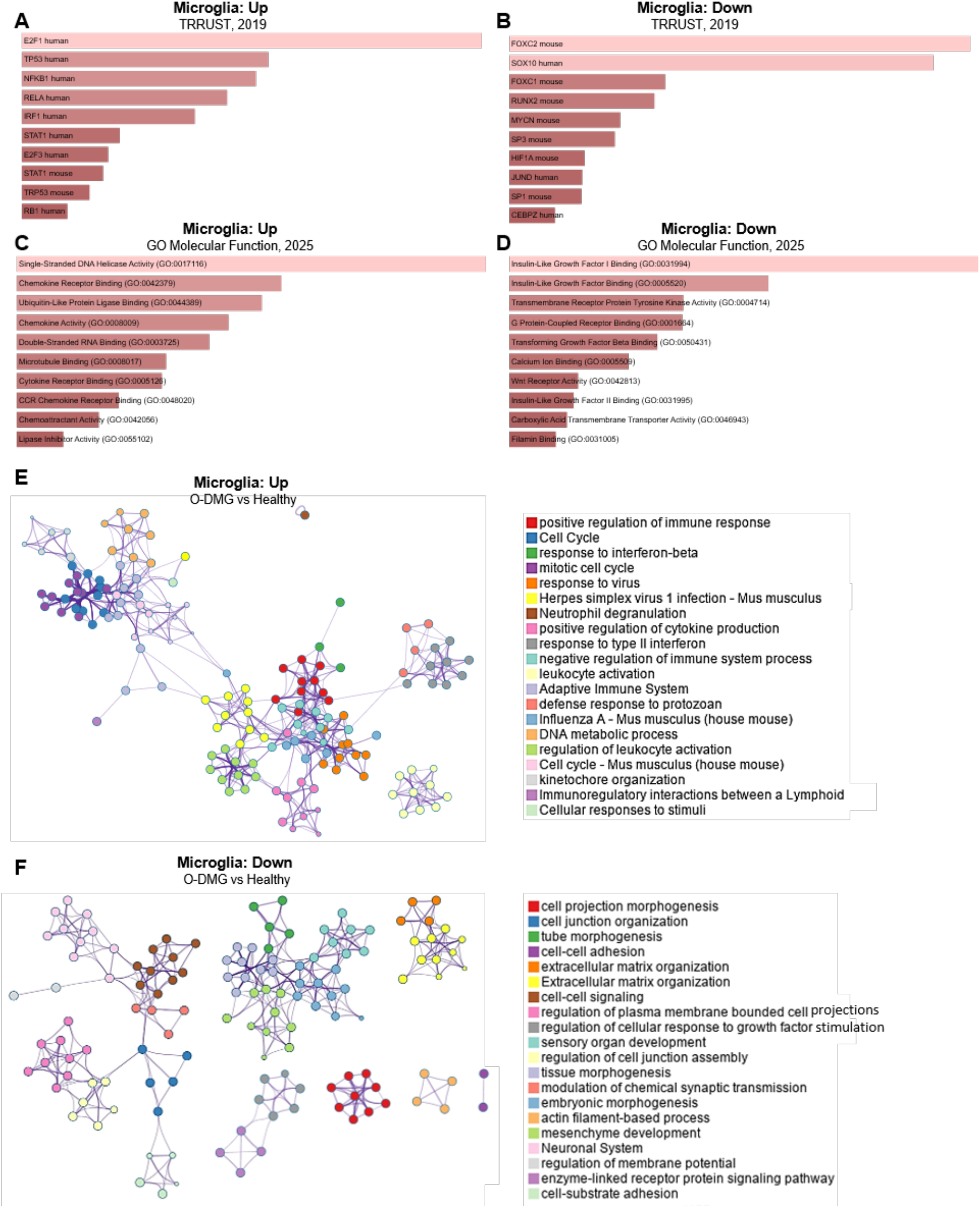
Microglia characterization in DMG tumours. Bulk RNA-sequencing of Microglia sorted from orthotopic (O-DMG) tumour region versus healthy hindbrain/midbrain (MB/HB): Gene enrichment analysis with Enrichr showing upregulated and downregulated: **A-B**) TTRUST transcription factor network analysis; **C-D**) GO Molecular function; **E-F**) Metascape enrichment network showing relationships among significantly enriched upregulated and downregulated pathways; n=3/group.

**Fig. S9:**
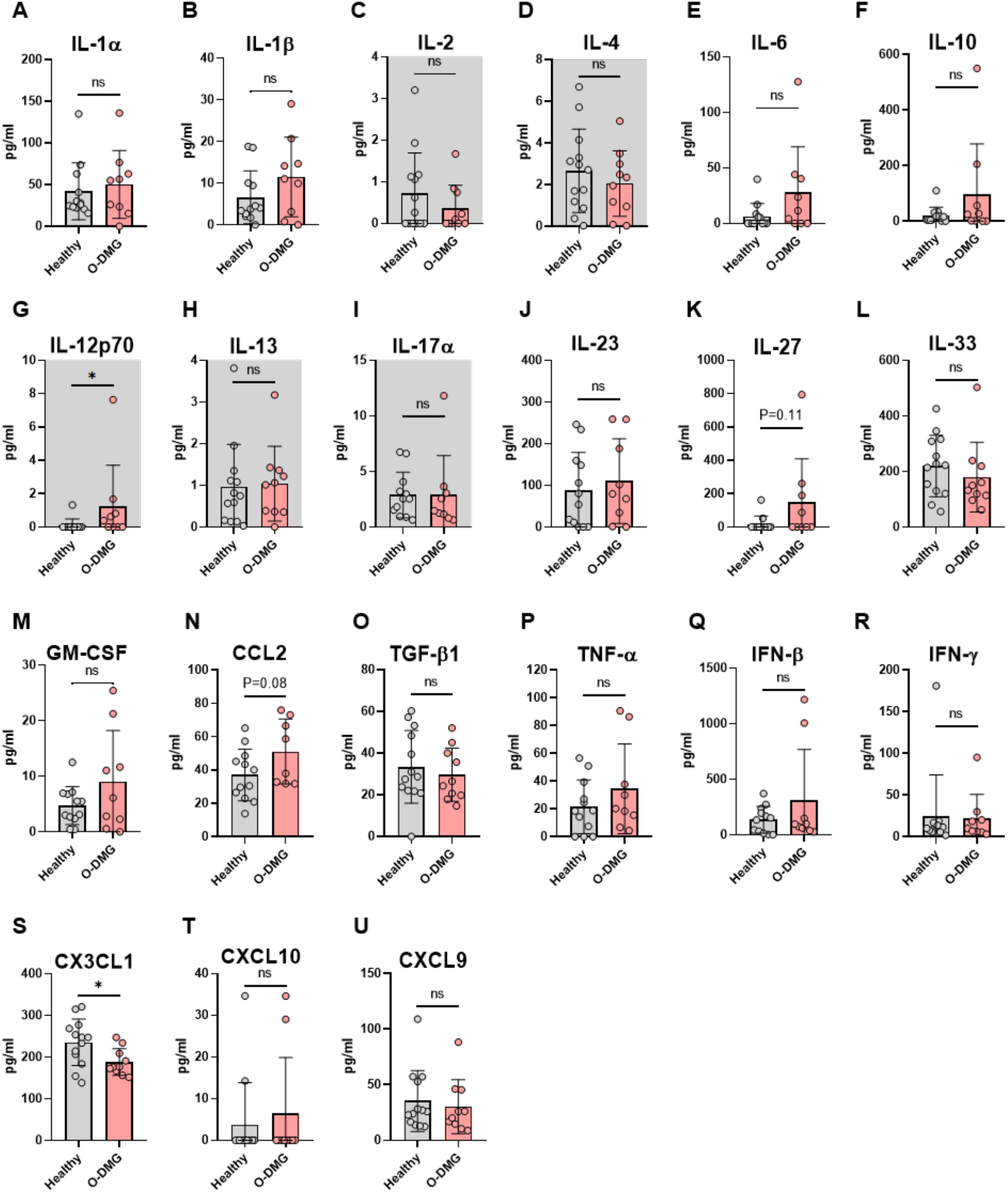
Cytokine quantification in healthy versus orthotopic (O-DMG) tumour-bearing C57BL/6 mice. LEGENDplex assay was performed to determine cytokines levels in blood plasma: **A**) IL-1α; **B**) IL-1β; **C**) IL-2; **D**) IL-4; **E**) IL-6; **F**) IL-10; **G**) IL-12p70; **H**) IL-13; **I**) IL-17α; **J**) IL-23; **K**) IL-27; **L**) IL-33; **M**) GM-CSF; **N**) CCL2; **O**) TGF-1β; **P**) TNF-α; **Q**) IFN-β; **R**) IFN-γ; **S**) CX3CL1; **T**) CXCL10; **U**) CXCL9. Cytokines below 5-15 pg/ml detection threshold were coloured with grey background. n=8-13/group; ns – non-significant, *p<0.05.

**Fig. S10:**
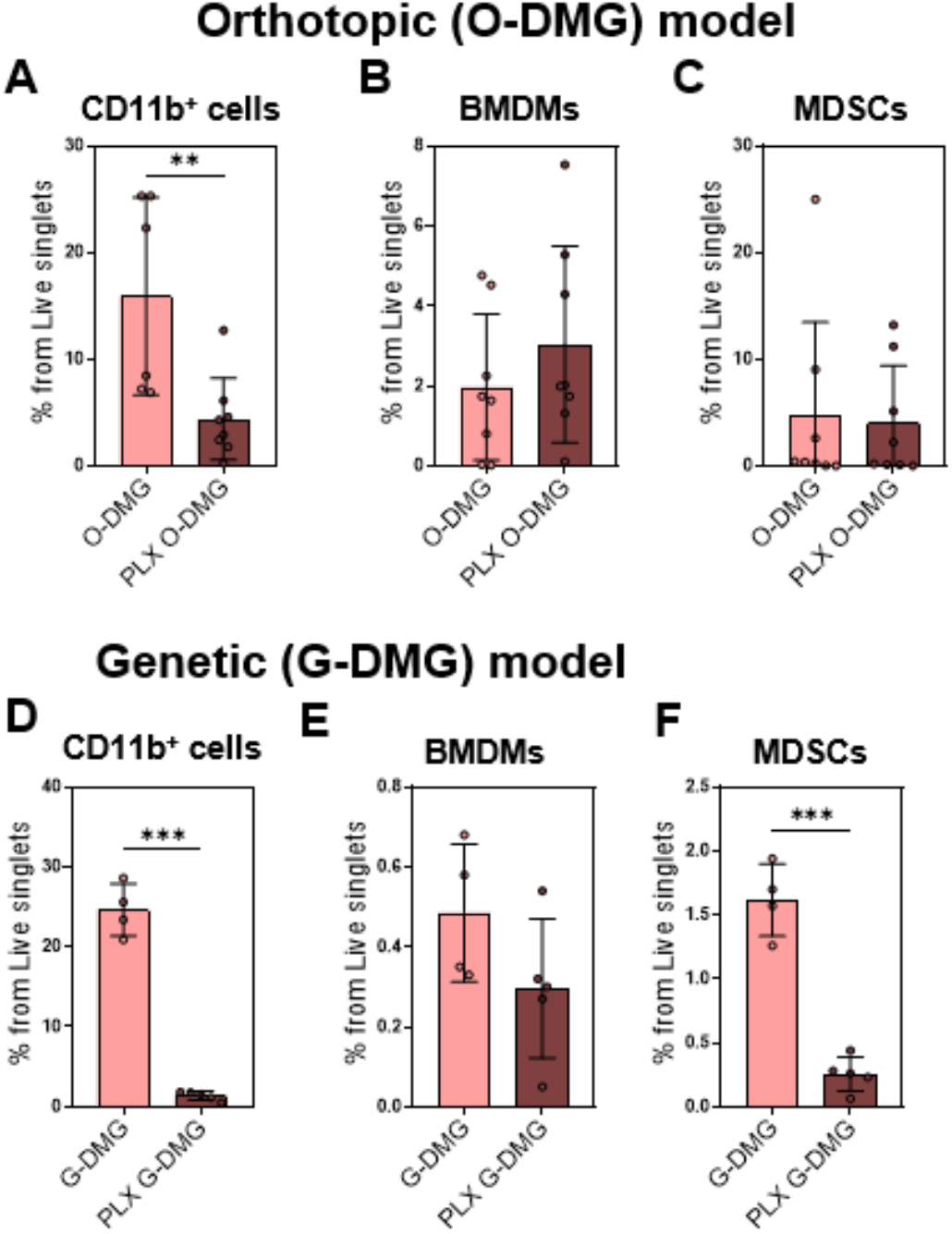
PLX3397 treatment alters abundances of CD11b^+^ myeloid cells, macrophages and MDSCs in the orthotopic (O-DMG) and genetic (G-DMG) tumour models. Flow cytometry analysis of: A) CD11b^+^ in O-DMG; B) CD11b^+^ in G-DMG; C) Macrophages (CD11b^+^CD49d^+^P2RY12^-^) in O-DMG tumour region; D) Macrophages in G-DMG; E) MDSCs (CD11b^+^CD49d^int^P2RY12^int^) in O-DMG; F) MDSCs in G-DMG; n=4-8/group. Significance: *p<0.05, **p<0.005, ***p<0.001, otherwise non-significant.

**Fig. S11:**
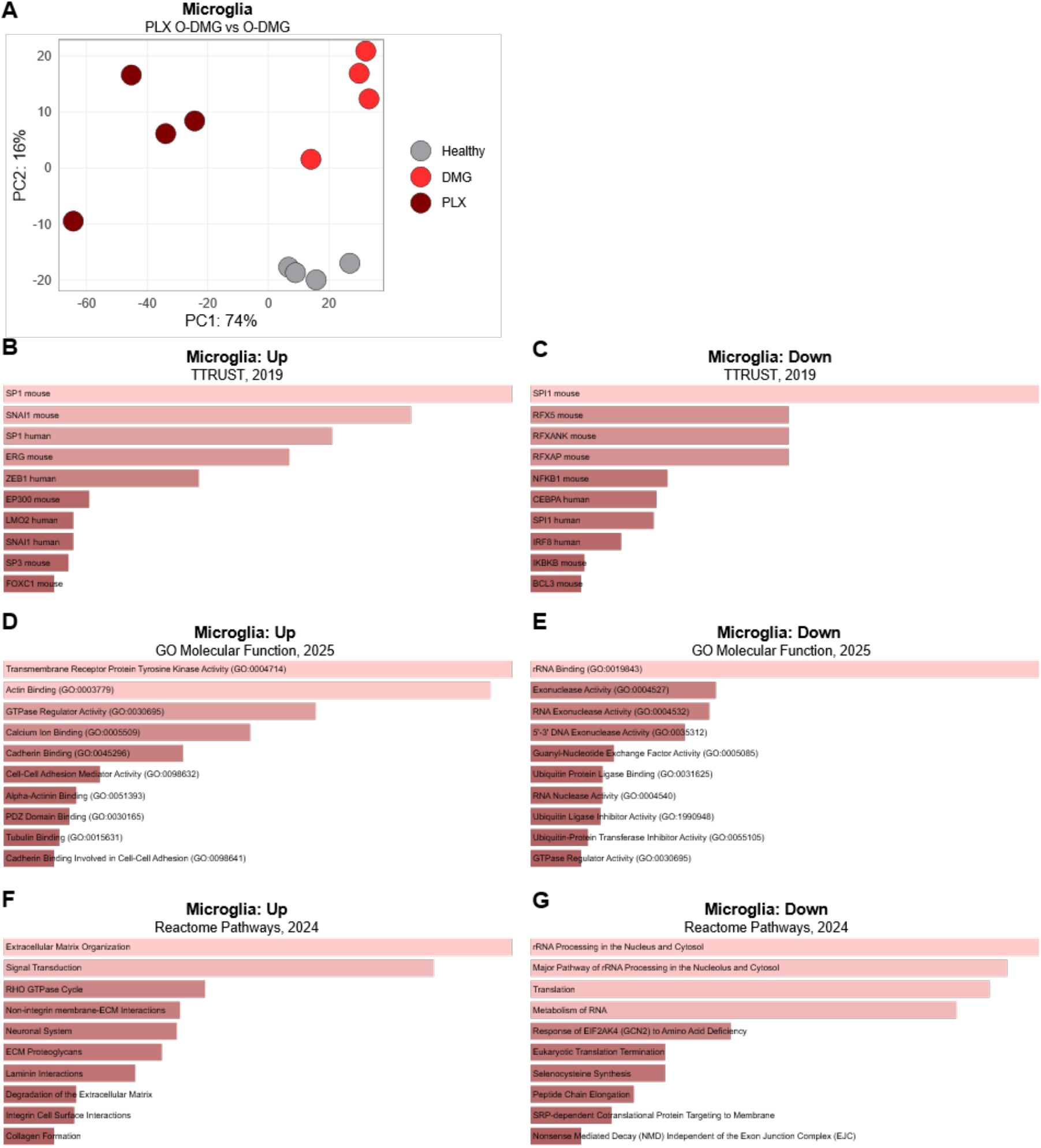
Microglia characterization in orthotopic (O-DMG) tumours after PLX3397 treatment. Bulk RNA-sequencing of Microglia (CD11b^+^CD49d^-^P2RY12^+^) sorted from untreated versus PLX3397-treated O-DMG tumours: **A)** Principal component analysis (PCA); Gene enrichment analysis with Enrichr showing upregulated and downregulated: **B-C**) TTRUST transcription factor network; **D-E**) GO Molecular function; **F-G**) Reactome pathways; n=4/group.

**Fig. S12:**
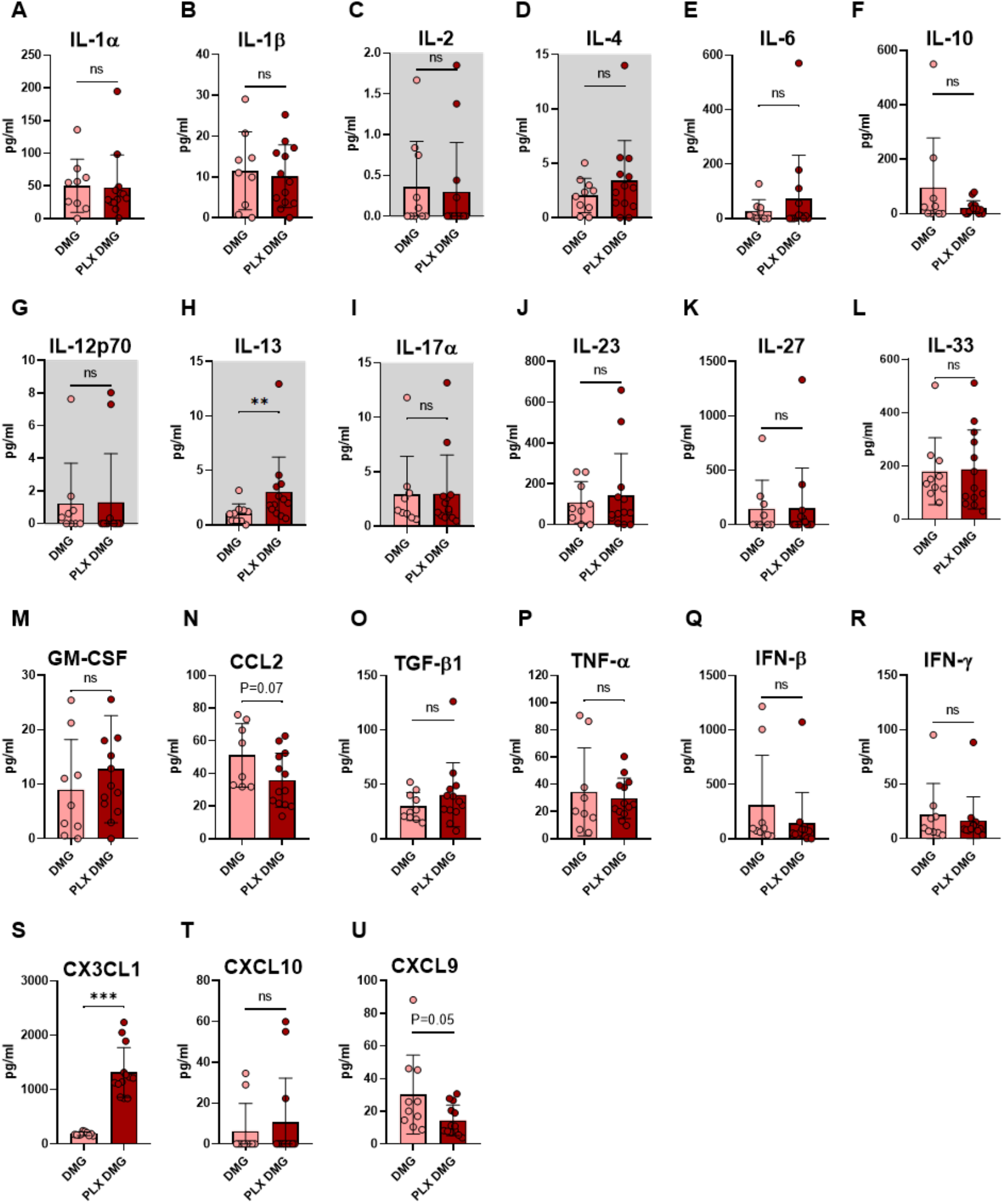
Cytokine quantification in untreated versus PLX3397-treated O-DMG tumour-bearing C57BL/6 mice. LEGENDplex assay was performed to determine the amount of cytokines in blood plasma: **A**) IL-1α; **B**) IL-1β; **C**) IL-2; **D**) IL-4; **E**) IL-6; **F**) IL-10; **G**) IL-12p70; **H**) IL-13; **I**) IL-17α; **J**) IL-23; **K**) IL-27; **L**) IL-33; **M**) GM-CSF; **N**) CCL2; **O**) TGF-1β; **P**) TNF-α; **Q**) IFN-β; **R**) IFN-γ; **S**) CX3CL1; **T**) CXCL10; **U**) CXCL9. Cytokines below 5-15 pg/ml detection threshold were coloured in grey background. n=9-13/group; ns – non-significant, **p<0.005, ***p<0.001.

**Fig. S13:**
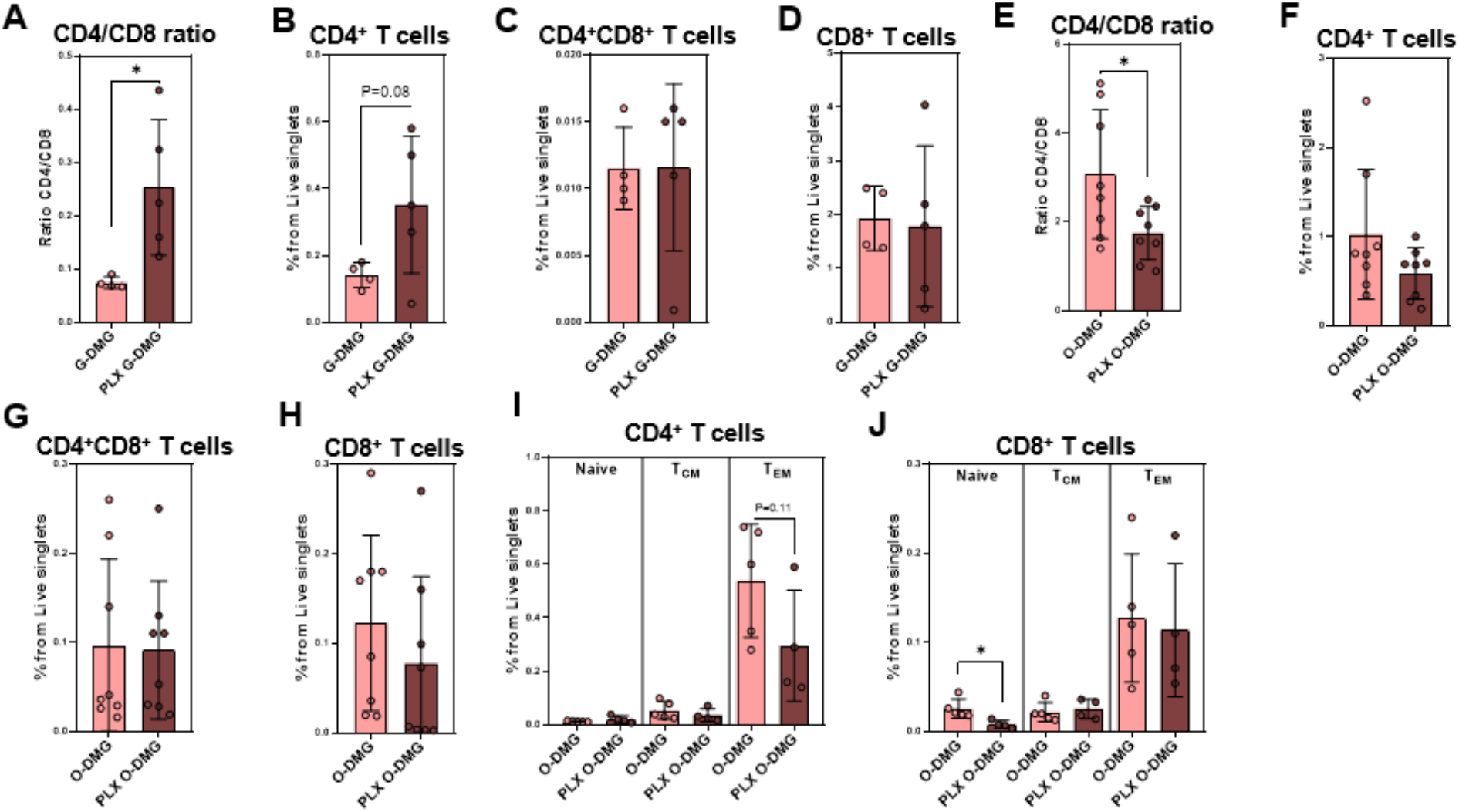
CD4^+^ T cell characterization in DMG tumours after PLX3397 treatment. Flow cytometry analysis of: **A**) CD4 to CD8 ratio in untreated versus PLX3397-treated G-DMG tumour region. The percentage of **B**) CD4^+^ T cells and **C**) CD4^+^CD8^+^ double-positive and **D**) CD8^+^ T cells from live singlets in untreated and PLX3397-treated G-DMG tumour region. **E**) CD4 to CD8 ratio in untreated versus PLX3397-treated O-DMG tumour region. The percentage of **F**) CD4^+^ T cells and **G**) CD4^+^CD8^+^ double-positive and **H**) CD8^+^ T cells from live singlets in untreated and PLX3397-treated O-DMG tumour region; n=4-8/group. Significance: *p<0.05, **p<0.005, otherwise non-significant. Naive, central-memory (T_CM_) and effector memory (T_EM_) subsets in O-DMG model in **I**) CD4^+^ T cells and **J**) CD8^+^ T cells.

**Fig. S14:**
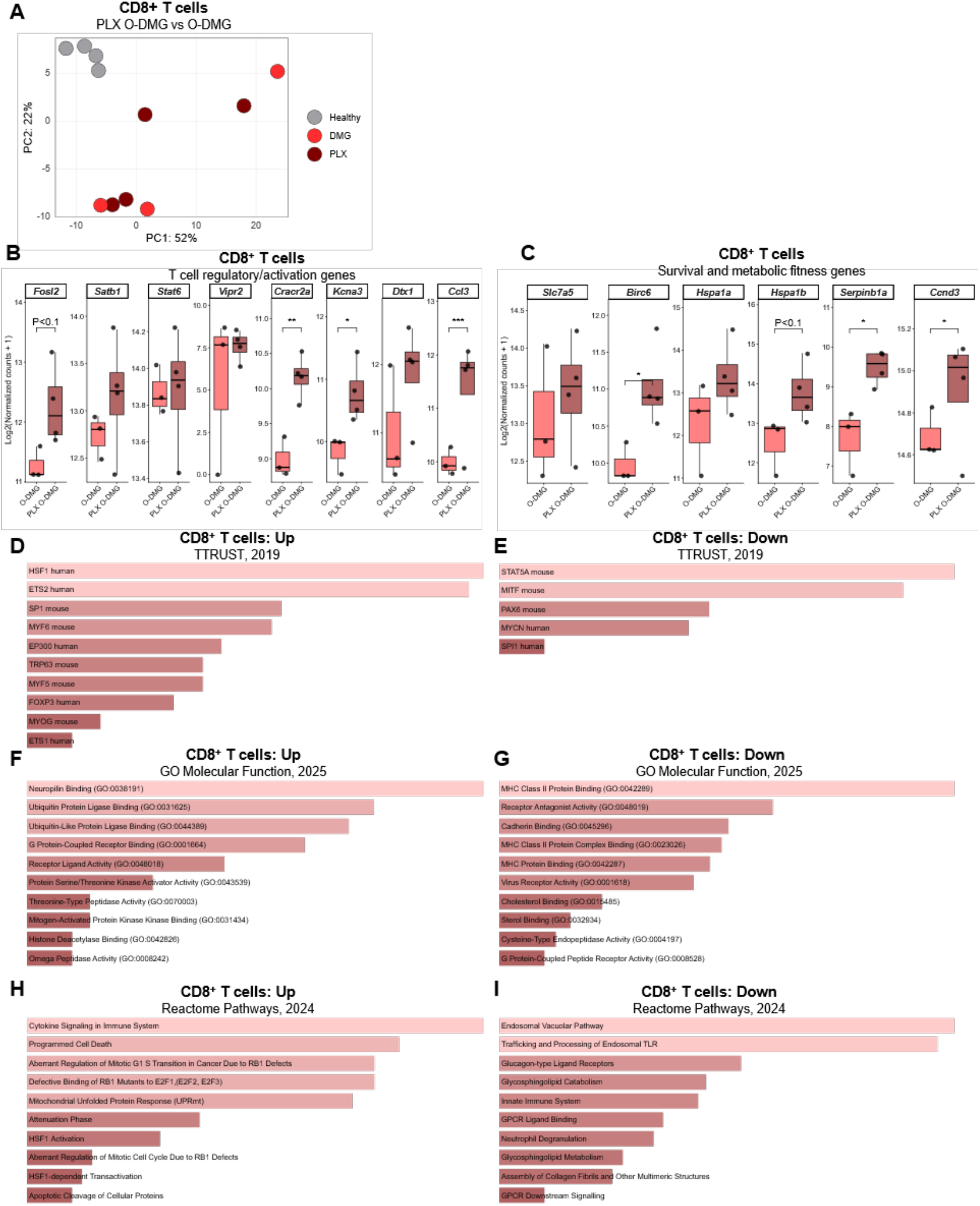
CD8^+^ T cell characterization in DMG tumours treated with PLX3397. Bulk RNA-sequencing of CD8^+^ T cells sorted from untreated versus PLX3397-treated O-DMG tumours. **A**) principal component analysis (PCA). Normalized transcript abundance of: **B**) Effector T cell regulatory/activation genes; **C**) T cell survival and metabolic fitness genes. Gene enrichment analysis with Enrichr showing upregulated and downregulated: **D-E**) TRRUST transcription factor network; **F-G**) GO Molecular function; **H-I**) Reactome Pathways; n=3-4/group; Significance: *p<0.05, **p<0.005, ***p<0.001, otherwise non-significant.

**Fig. S15:**
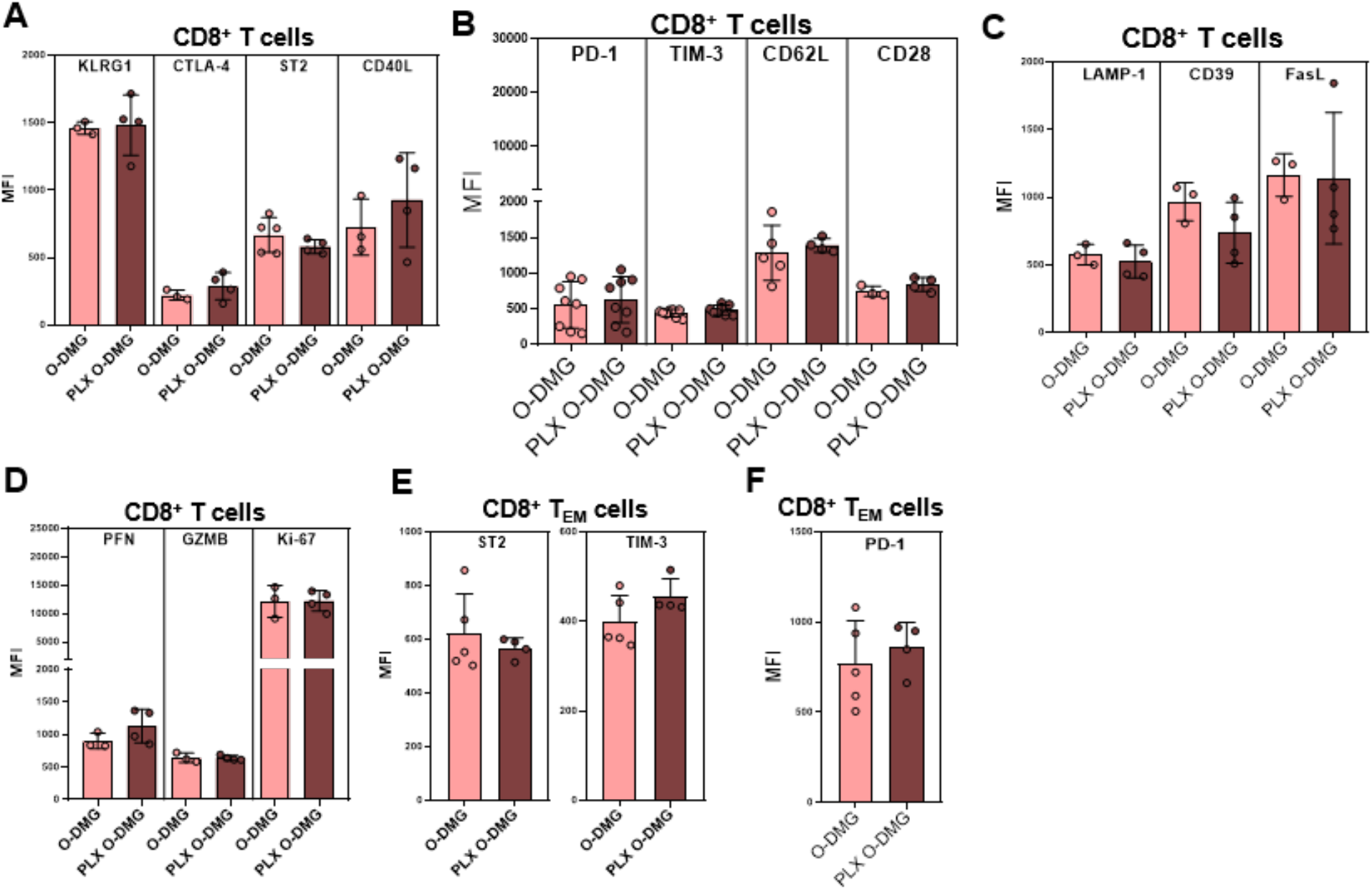
Flow cytometry phenotyping of CD8^+^ T cells in untreated versus PLX-treated O-DMG tumours. **A**) Median fluorescence intensity (MFI) of KLRG1, CTLA-4, ST2, CD40L; **B**) PD-1, TIM-3, CD62L, CD28; **C**) LAMP-1, CD39, FasL; **D**) perforin (PFN), granzyme B (GZMB) and Ki-67 from CD4^+^ T cells. **E**) MFI of ST2, TIM-3 and **F**) PD-1 from CD4^+^ T effector-memory cells (T_EM_; CD62L^-^CD44^+^); n=3-8/group. Significance: *p<0.05, **p<0.005, otherwise non-significant.

**Fig. S16:**
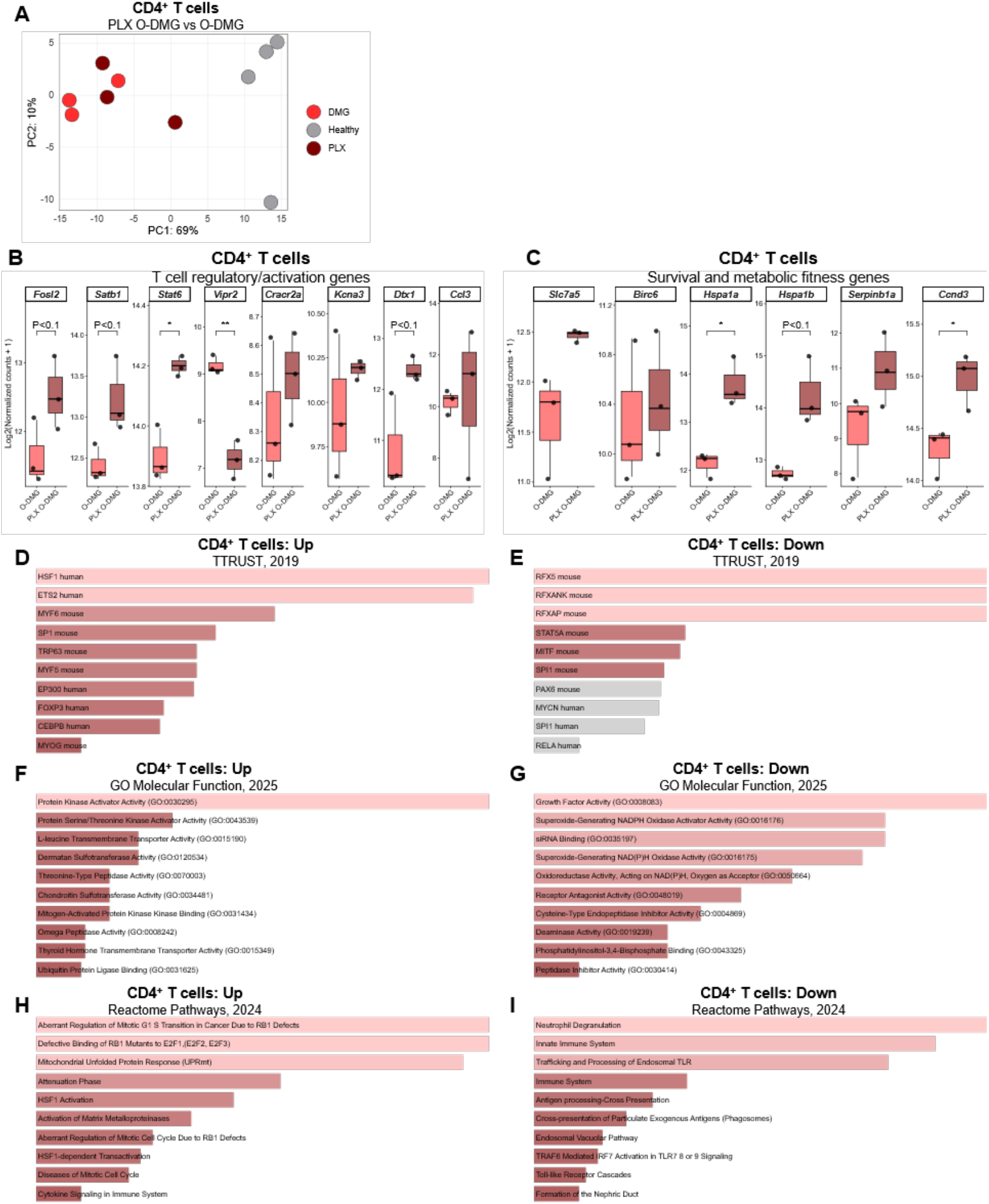
CD8^+^ T cell characterization in DMG tumours treated with PLX3397. Bulk RNA-sequencing of CD4^+^ T cells sorted from untreated versus PLX3397-treated O-DMG tumours. **A**) principal component analysis (PCA). Normalized transcript abundance of: **B**) Effector T cell regulatory/activation genes; **C**) T cell survival and metabolic fitness genes. Gene enrichment analysis with Enrichr showing upregulated and downregulated: **D-E**) TRRUST transcription factor network; **F-G**) GO Molecular function; **H-I**) Reactome Pathways; n=3-4/group; Significance: *p<0.05, **p<0.005, ***p<0.001, otherwise non-significant.

**Fig. S17:**
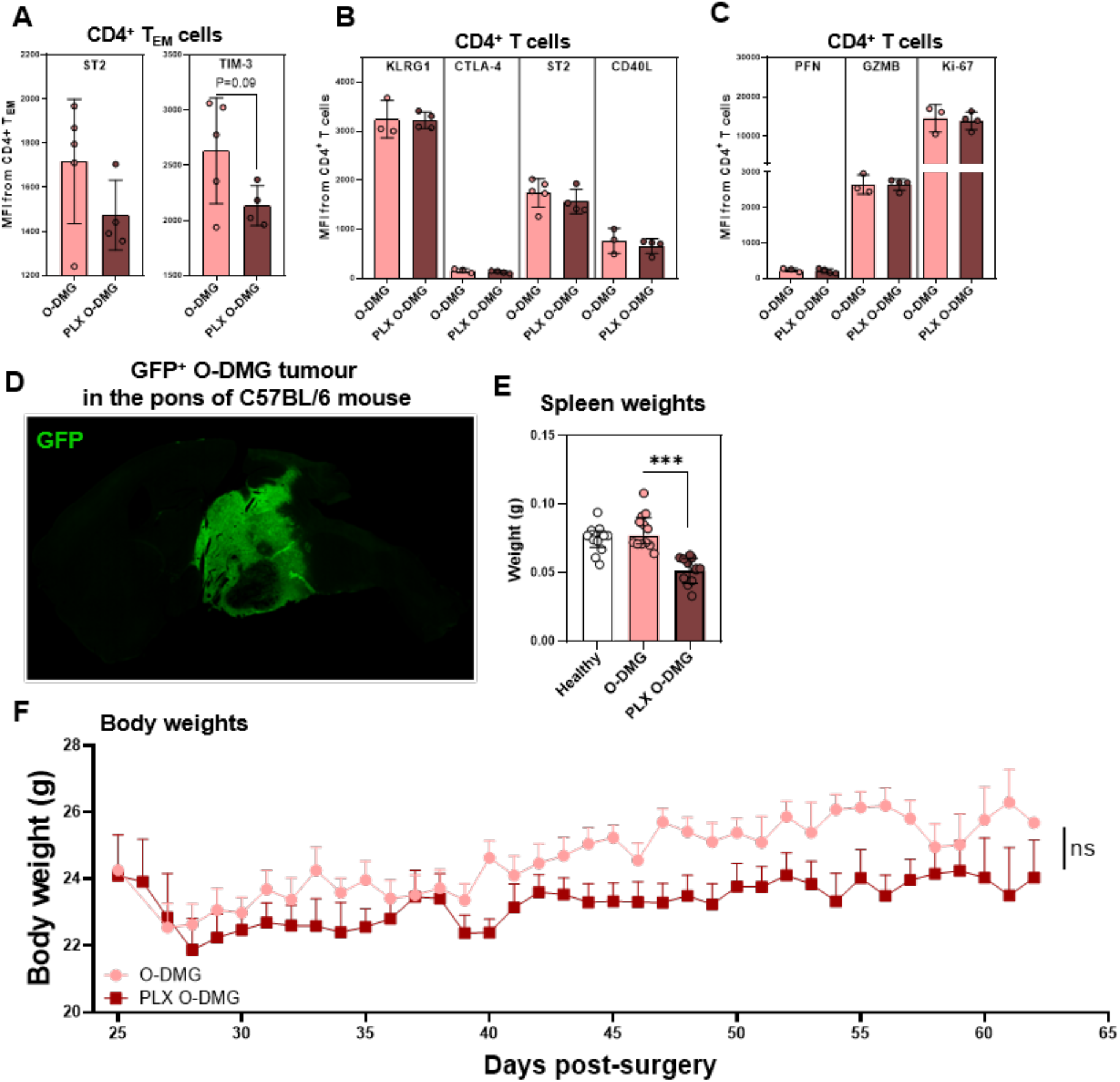
Flow cytometry phenotyping of CD4^+^ T cells in untreated versus PLX-treated O-DMG tumours and impact of PLX3397 treatment on the C57BL6 mice survival and tumour burden. Flow cytometry of CD4+ T cells in untreated and PLX-treated O-DMG tumours: A) Median fluorescence intensity (MFI) of ST2 and TIM-3 from CD4^+^ T effector-memory cells (T_EM_; CD62L^-^CD44^+^). B) MFI of KLRG1, CTLA-4, ST2, CD40L from global CD4^+^ T cells. **C**) perforin (PFN), granzyme B (GZMB) and Ki-67 from global CD4^+^ T cells; n=3-8/group. Significance: *p<0.05, **p<0.005, otherwise non-significant. **D**) Immunofluorescence microscopy with Olympus slide scanner of untreated GFP+ O-DMG tissue to confirm tumour localization (10X magnification). Comparisons between untreated and PLX3397-treated O-DMG tumour-bearing mice: **E**) Spleen weights, n=12-13/group. **F**) Body weight, n=18-19/group.

